# Constrained actin dynamics emerges from variable compositions of actin regulatory protein complexes

**DOI:** 10.1101/525725

**Authors:** Ulrich Dobramysl, Iris K Jarsch, Hanae Shimo, Yoshiko Inoue, Benjamin Richier, Jonathan R Gadsby, Julia Mason, Astrid Walrant, Richard Butler, Edouard Hannezo, Benjamin D Simons, Jennifer L Gallop

**Affiliations:** Gurdon Institute, University of Cambridge, Cambridge, CB2 1QN, UK; Department of Biochemistry, University of Cambridge, UK, CB2 1GA, UK; Institute of Science and Technology Austria, 3400 Klosterneuburg, Austria; Department of Applied Mathematics and Theoretical Physics, University of Cambridge, Cambridge, CB3 0WA, UK

## Abstract

Assemblies of actin and its regulators underlie the dynamic morphology of all eukaryotic cells. To begin to understand how diverse regulatory proteins work together to generate actin-rich structures we tracked the assembly of actin regulators and their relative proportions in a cell-free system that generates filopodia-like structures (FLS). We found that heterogeneous mixtures of regulators could give rise to morphologically similar structures and that the FLS actin bundles exhibited simple dynamic behaviour of growth and shrinkage. To explain these observations, we combined experiment with theory, and found that stochastic fluctuations between redundant actin regulatory subcomplexes can account for the actin dynamics. Comparing the localizations of a variety of endogenous actin regulators in *Drosophila* embryos and distributions of filopodia lengths yielded similar conclusions of heterogenous actin regulatory complexes and filopodia lengths governed by a stochastic growth process. Our results explain how weakly-associating assemblies of regulatory proteins can produce robust functional outcomes.

## Introduction

The regulation of actin polymerization is crucial for numerous cell functions, including cell migration, adhesion and epithelial closure (1, 2) and is often disrupted in disease, such as cancer metastasis and intracellular infection by pathogens (3–6). Micron-scale structures of actin filaments and associated regulators provide the mechanical infrastructure for the cell via transient membrane-bound complexes that somehow orchestrate large scale cytoskeletal remodelling (7, 8). Filopodia, with their characteristic membrane-associated “tip complex” where new actin monomers are incorporated, are one such example (9, 10). Current models for filopodia formation posit key roles for either (1) formins in *de novo* nucleation processes (11–13), (2) a preexisting actin network generated by the Arp2/3 complex and bundled by fascin (14–16) or (3) the clustering and outward projection of membrane-bound SH3 domain containing adaptor proteins that recruit Enabled (Ena)/ Vasodilator-stimulated phosphoprotein (VASP) proteins coupled to the wider actin machinery (17–19). One way to reconcile these models is to postulate the existence of subtypes of filopodia based on their mechanism of formation (20–24). However, a unifying picture is still lacking, such that the nature, number and dynamics of putative filopodia subtypes remains in question.

Purified reconstituted systems using membranes, several signaling proteins and receptors have revealed that a semi-dynamic network of actin regulators can form from multivalent interactions, and that these can determine the locations of actin filament assembly (25–27). While we now appreciate how such condensates can be generated by their underlying molecular characteristics, open questions remain. In particular, how the molecular nature and composition of the regulatory protein assemblies contribute to their functional output in producing large scale actin structures remains unknown (8, 28). While some actin regulatory subcomplexes are known to have defined stoichiometry (for example the Wiskott-Aldrich syndrome protein family Verprolin-homologous protein (WAVE) complex (29)), others such as neural Wiskott Aldrich syndrome protein (N-WASP) can participate at variable stoichiometries and be highly dynamic (25, 30). The complex of actin regulators such as those found at filopodia tips, presumably lies somewhere between a defined complex and a loose association of signaling and effector proteins. We do not know whether specific combinations and stoichiometries of actin bundlers, nucleators, and elongators are needed to create an actin bundle and if there are, what these ratios would be.

We previously described a cell-free system comprised of *Xenopus* egg extracts and a phosphatidylinositol (4,5)-bisphosphate supported lipid bilayer that grows actin bundles from a membrane-bound complex of actin regulators that is highly amenable to quantitative microscopy (Fig. 1A) (19). Their compositional similarity to filopodia, and their ability to grow many microns long, led us to call them filopodia-like structures (FLS). Although the complex of actin regulators is at the base of the FLS, in filopodia the complex of actin regulators is found at the tip; therefore, we refer to the complex of actin regulators in FLS as the tip complex by analogy with filopodia. Both the tip complex and the actin bundle can be readily followed by timelapse fluorescence microscopy at the bilayer surface and up from it, in the z axis. FLS are not a strict filopodia mimic as membrane does not surround the shaft, and N-WASP is employed as an Arp2/3 complex activator rather than the closely related Scar/WAVE protein that may be fulfilling this role in cells. Nonetheless, FLS contain bundled actin filaments and new actin monomers are incorporated at a membrane-localized complex that includes Ena, VASP and Diaphanous-related formin 3 (Diaph3). The actin bundling protein, fascin is also present throughout the shaft.

**Fig. 1.**
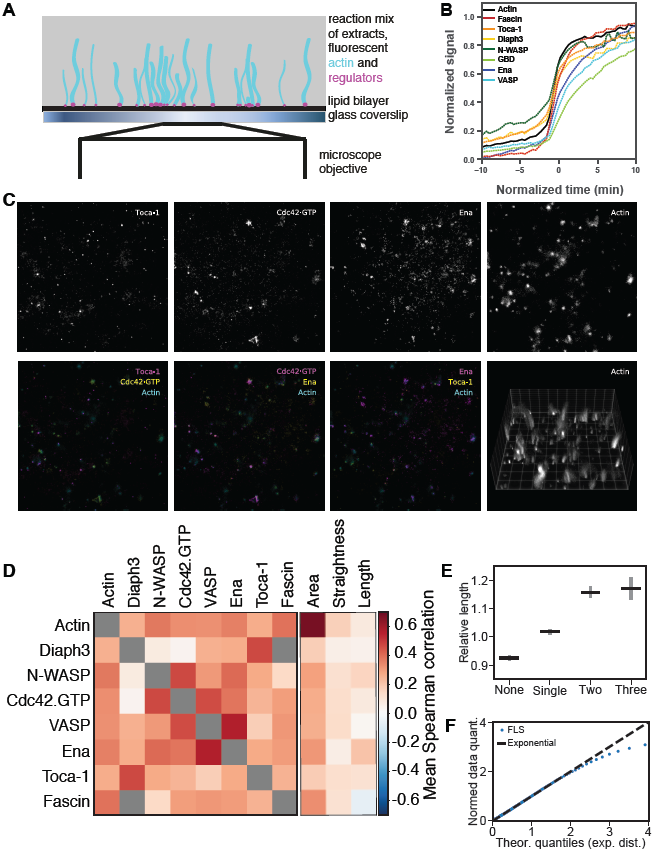
Actin regulatory proteins are heterogeneous and redundant and exhibit preferred subcomplexes. (A) Schematic diagram of the FLS assay. (B) Accumulation of actin regulators over time as FLS form. Actin n = 19974, fascin n = 2898, TOCA-1 n = 19893, Diaph3 n = 3811, N-WASP = 1743, GBD n = 7839, Ena n = 1828, VASP n = 1857. (C) Complexes of diverse stoichiometry generate filopodia-like structures (FLS) *in vitro*. Example of three different actin regulatory proteins (TOCA-1, Cdc42.GTP and Ena) at the membrane together with a 3D reconstruction of z-stack of the actin contained in FLS growing from a supported bilayer. Note the diversity of regulatory protein combinations observed under the different actin structures (coloured images in the bottom row). (D) Matrix showing Spearman protein-protein intensity correlation and morphology-protein intensity correlation values calculated separately per field of view and subsequently averaged. See figure supplement for N and n numbers. Gray boxes mean exact correspondence (Diaph3 and fascin are both GFP tagged). (E) Relative change in FLS length for FLS with zero, one, and any two or three observed proteins enriched compared to the total population length mean, calculated for each field of view and subsequently averaged. Error bars indicate the 95% confidence interval of the mean. (F) Quantile-quantile plot showing that FLS lengths are exponentially distributed over a wide range. The line tails off when the n becomes low.

Here we tracked the assembly of the actin regulators to the FLS tip complex in time, as well as their intensities in relation to the actin bundle. We found that the tip complex assembles cooperatively in an F-actin dependent manner, that the final complex is heterogeneous in composition, and that proteins can turn over within it. Despite their heterogeneity of accumulation, we find that some of the actin regulators comprising the tip complex form subcomplexes. However, the balance of assembly and disassembly of the actin gives rise to bundle morphologies that are independent of the precise stoichiometry and composition of the regulators within the tip complex. This observation suggests a redundancy among actin regulators and is puzzling because one would naively expect the concentration of protein to be directly linked to morphological properties, such as length. To establish the link between FLS morphology and the heterogeneity in actin regulator composition we employed theoretical modelling. This has revealed a simple and universal law for the redundant assembly and growth dynamics of actin structures via the action of independent subcomplexes of actin regulators. Moreover, by using fluorescent protein knock-in and measurements of filopodia in *Drosophila* embryos we could show similar heterogeneity in the tip complexes and length characteristics of filopodia *in vivo*. Our work reveals how a dynamic and redundant actin regulatory complex gives rise to morphological and behavioural robustness of the resulting actin bundles.

## Results

### Heterogeneity in actin regulators during FLS assembly

To track the assembly of the tip complex of the FLS, for example to determine whether there is a distinction between actin regulators that nucleate actin filaments, and those that bundle or extend actin filaments, we measured the intensites of actin regulatory proteins with time as FLS form (Fig. 1A,B). To add fluorescently labelled proteins into the extracts, we expressed and purified *Xenopus laevis* or *tropicalis* Transducer of Cdc42 activation-1 (TOCA-1), the G-protein binding domain of N-WASP to monitor Cdc42•GTP (GBD), N-WASP, Ena, VASP, Diaph3 and fascin (domain structures are shown in Figure 1 supplement 1A, gels in B). TOCA-1, Ena and N-WASP were expressed as SNAP-fusion proteins and chemically labelled with AlexaFluor (AF) 647 and AF 488. VASP was engineered to contain a cysteine close to the N-terminus for labelling with AF 568 and AF 647 maleimide, as previously published (31). Fascin and Diaph3 were purified as N-terminal GFP-fusion proteins, and GBD as an N-terminal fusion protein with red fluorescent protein pmKate2. To assess the concentration of labelled protein added relative to the endogenous protein provided by the extracts, we measured the concentrations of TOCA-1, Ena, VASP, N-WASP and fascin in the high-speed supernatant extracts using quantitative western blotting (Fig. 1 supplement 1C-E). We obtained similar results to quantitative proteomics of *Xenopus laevis* egg cytoplasm (32). We optimized our microscopy to add as little recombinant protein as possible while retaining sufficient signal-to-noise ratio in the images. The resulting reaction mix contained proteins at 1:1-1:2 labelled:unlabelled ratio. To monitor the simultaneous accumulation of different actin regulators to the FLS tip complex together with elongation of the actin bundle, we used highly inclined and laminated optical sheet (HILO) illumination in up to 3 wavelengths and widefield illumination or spinning disc confocal optical sectioning in 3D. We employed rapid sequential timelapse imaging of actin and other proteins in different combinations to visualize bundle dynamics. We developed an ImageJ plugin, “FLS Ace” that segments individual 3D actin structures from the confocal or widefield channel. Furthermore, it quantifies intensities from any additional HILO channels at the membrane underneath. This enabled us to extract protein intensities at the tip complex and along the shaft of actin bundles from the acquired sets of images. Linear assignment between timepoints allows tracking of single FLS throughout their lifetime, as well as the detection of actin regulators that accumulate at the membrane prior to actin polymerization (Fig. 1 supplement 2).

These experiments revealed that the initiation of new FLSs occurs throughout the experiment with a peak at 3 min (Fig. 1 supplement 3A). To be able to compare individual FLSs irrespective of their time of initiation, we defined the timepoint at which a structure was first detected as t0 in “FLS-time” as compared to “experiment-time”. While an FLS that appears later in the experiment can reach the same length as those initiated early, the size of their tip complex area is on average slightly smaller, probably due to limitations of either a protein component in the reaction mixture or physical space on the membrane (Fig. 1 supplement 3B,C). Normalizing the data to FLS-time, we can see a cooperative assembly of the actin regulators to the tip complex, where the largest increase in fluorescence with time occurs at the same time as peak actin accumulation just prior to t0, defined as the first detection of a 3D structure (Fig. 1B). The recruitment of the Cdc42•GTP probe, GBD, was retarded relative to the other proteins, which is likely due to competition with endogenous N-WASP (Fig. 1B). While there is very little variation in the timing of protein accumulation, considerable variability in the mean intensity at steady-state is evident in the data (Fig. 1 supplement 3D). To test whether the assembly of the complex is driven by F-actin (the central concept underlying the convergent elongation model of filopodia formation), we tested which of our proteins still assembled in the presence of actin monomer sequesting drug, Latrunculin B (LatB) to prevent the polymerization of actin into filaments. As observed previously, our experiments showed that TOCA-1 was still present, though is somewhat reduced, in the absence of F-actin (19), with comparable reductions of Ena and N-WASP (Fig. 1 supplement 4). The recruitment of fascin and VASP was prevented, similar to previous observations with Diaphanous 2 (19), and the presence of Cdc42•GTP at the membrane partially affected (Fig. 1 supplement 4). This is in accordance with an initial slow rise in membrane/Cdc42•GTP interactors TOCA-1 and N-WASP laying a foundation for the recruitment of actin and other regulatory proteins with cooperative assembly of the tip complex coinciding with the peak actin accumulation (Fig. 1B).

To assess the compositional heterogeneity of the final assembled FLS tip complex in more detail we took snapshot images 20 mins after starting the assay, including each protein in combination with the others, in double and triple combinations. Otherwise indistinguishable FLS actin bundles emerge from tip complexes whose composition differs both qualitatively and quantitatively (Fig. 1C shows example data from one combination). We looked at the correlation between each pair of actin regulators, and the correlation of each regulator with the actin intensity in the bundle (Fig. 1D). We found several protein pairs that exhibit strong intra-FLS correlations: Ena/VASP, TOCA-1/Diaph3, VASP/Cdc42•GTP and N-WASP/Cdc42•GTP (Fig. 2D, see Fig. 1 supplement 5 for number of FLS). The high correlation between Ena and VASP can be explained by their known ability to form heterotetramers (33) and VASP is also reported to cooperate with Cdc42 (17). Diaph3 is a SH3 domain binding partner of TOCA-1 previously observed by co-immunoprecipitations (34), and Cdc42•GTP is known to be a major input into N-WASP activation (35). Nearly all other pairs show weak positive correlations, with the exceptions of N-WASP/Diaph3 and Cdc42•GTP/Diaph3, which exhibit close to zero correlation i.e. they distribute randomly relative to each other (Fig. 1D).

**Fig. 2.**
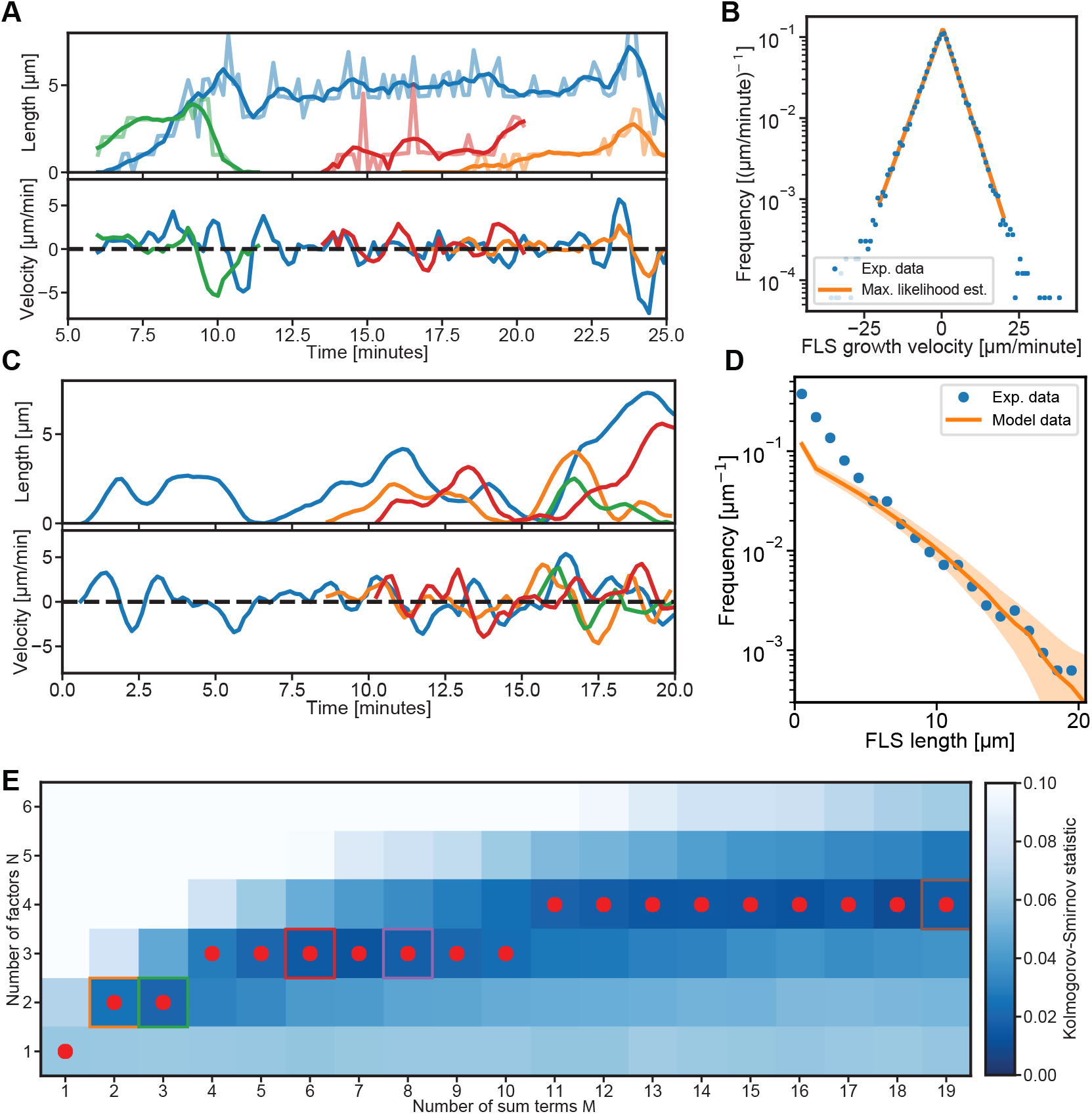
Mathematical model for FLS growth captures experimental FLS length distribution and growth dynamics. (A) Four smoothed example FLS length trajectories (top) and their instantaneous growth velocities (bottom). The colours are different examples. (B) Histogram of measured FLS growth velocities (n = 114816 velocity observations) from tracked FLS trajectories with Maximum Likelihood Estimation fit of a bi-exponential Laplace distribution. (C) Four randomly selected smoothed example trajectories (top) generated from our sum-product model with their instantaneous velocities (bottom). Based on our observations that new FLS are spawned at random times throughout the assay we started our model simulations at random initiation points in the time interval between zero and 20 minutes. (D) FLS length distribution (blue circles, N=3193 observations) and corresponding data from simulating 10,000 model trajectories (solid orange line, shaded area is the 95% confidence interval of the histogram bin means). Simulation data histogram was scaled by the ratio between the median FLS lengths larger than 5 *μ*m and the median simulated FLS lengths larger than 5 *μ*m to visualize agreement between theory and experiment. (E) Comparison of the experimental normalized growth velocity distributions to the model for different combinations of number of sum terms M and number of product factors N, using the Kolmogorov-Smirnov goodness-of-fit statistic. Darker blue indicates a better fit. The best fit N for any given M is highlighted by a red dot and follows an approximate square root dependence. The growth velocity distribution fits for data in coloured boxes is shown in Fig. 2 supplement 1.

Low levels of fluorescence can result from both nonspecific protein-protein interactions and by specific interactions between proteins of low abundance. The difference between these two possibilities is obscured at low intensities, hence we verified our conclusions about the heterogeneity of FLS tip complexes by focusing on high protein intensity levels. To this end, we considered an FLS to be enriched with a given protein if it falls within the brightest 30% in the respective channel. This threshold resulted from taking the top half of FLS as ranked by their brightness while excluding FLS with protein levels below background. From experiments in which we labelled three different tip complex proteins (any three, in different combinations) we observed that 36% of FLS are enriched in one of the three proteins, 14% are enriched for two out of the three and only 2% have high levels of all three of the proteins. 48% of FLS are not enriched for any of the three proteins (where the other regulators that are in the extracts and unlabelled are presumably present). Broad heterogeneity is therefore apparent even when selecting the most fluorescent FLS. Comparing the observed percentages of FLS that are enriched for two given proteins with the percentages expected from random associations only reveals the same relationships as the correlation analysis: there is a significant positive association for all combinations except N-WASP/Diaph3 and Cdc42•GTP/Diaph3, which distribute randomly (Fig. 1 supplement 5 shows the percentage of FLS with two enriched proteins overlapping together with p-values indicating the significance of the percentage compared to random association).

Thus we find a generally permissive but not totally promiscuous association between actin regulators. The lack of correlation of Diaph3 with Cdc42•GTP may be due to competition with the GBD domain probe. However, in that case N-WASP should also be competitive with GBD; instead it shows a high correlation (Fig. 1D). The overall positive correlation between most proteins suggests a cooperativity with more proteins joining the assembly the more that are there, which agrees with the sigmoidal shape of protein assembly in the kinetic analysis (Fig. 1B). While some components are found to associate more frequently with each other, these data rule out the presence of discrete subtypes of FLS tip complex as in that scenario there would have been both strong positive and strong negative relationships.

We next examined the correlations between tip complex composition and FLS morphology (Fig. 1D). There was a low positive correlation between each of the regulators with the area of the complex, measured with fluorescent actin (Fig. 1D). FLS are somewhat wavy, though less so than the trajectories of *Listeria* comets in *Xenopus* egg extracts (36), however there was little correlation between FLS straightness and protein intensity (Fig. 1D). Indeed, there was also little correlation of any single protein intensity parameter with FLS length (Fig. 1D). This is surprising as one would expect there to be a strong role for some regulators to determine bundle lengths. FLS simultaneously enriched with two or three of the observed regulators were slightly, but significantly, longer than those with none or one (Fig. 1E, error bars show the 95% confidence interval). To determine whether we could discern any specifically short or long subtypes of FLS, which would be indicated by multimodal distributions, we plotted the length distributions. Instead, we found that the FLS lengths lay on a continuous exponential distribution (Fig. 1F shows a quantile:quantile plot comparing the data to a theoretical exponential distribution). Rather than a complex relationship between actin regulator compositions and lengths of FLS suggested by the diverse compositions of actin regulators, the exponential distribution points to a simple law governing FLS growth that is independent of the concentrations of actin regulators. Exponential distributions of lengths suggests a process dominated by stochastic incorporation of actin at functionally equivalent yet molecularly diverse tip complexes.

### FLS growth dynamics and theoretical framework

To draw a connection between the heterogeneity of actin regulators inside FLS tip complexes and the observed exponential length distribution, we developed a theoretical framework to describe FLS length dynamics. We applied this to the lengths of large numbers (>100,000) of individual FLS measured with high time resolution (Fig. 2A, B). By analogy to a study of lamellipodia dynamics (37), after an initial rapid growth phase, the FLS length cycles between elongation and retraction phases (Video 1, Fig. 2A). From these data, we calculated histograms of growth velocities, which had exponential tails both in positive (growth) and negative (shrinkage) directions and were largely symmetric. This bi-exponential shape is a Laplace distribution, which fits the data to a remarkable degree (Fig. 2B), unlike others such as a normal or power-law distributions. This is especially striking because the Laplace distribution is controlled only by a single parameter (its variance).

We next sought to understand the simplicity of such a growth distribution apparently arising from the complex networks of molecular associations observed between the actin regulatory proteins at the tip complex. Theoretically, a Laplace distribution can emerge when four normally distributed random variables *X*_1_,*X*_2_,*X*_3_ and *X*_4_ are combined as the sum of products *V* = *X*_1_*X*_2_ + *X*_3_*X*_4_ (see supplementary information for details). We are interpreting the variable *V* as the instantaneous growth velocity and hypothesized that normally distributed X’s could be caused by stochastic (Gaussian) fluctuations of actin regulatory proteins. Product terms can arise from complex-forming molecular reactions *X*_1_ + *X*_2_ → *X*_1_*X*_2_, with the complex *X*_1_*X*_2_ producing FLS elongation or shrinkage, while the sums represent the independent effect of two complexes *X*_1_*X*_2_ and *X*_3_*X*_4_ on velocity. When fluctuations are large compared to baseline concentrations, this provides a simple framework in which Laplace distributions can emerge from stochastic protein dynamics (see supplementary information for details). The dynamics of the concentration fluctuations X are controlled by a single parameter, the relaxation rate *θ*, which determines how quickly a concentration returns to its baseline after a random change. Based on this model, we could simulate velocity trajectories and, via numerical integration, FLS length trajectories (Fig. 2C). We collected length data points at the 20-minute time point, both in our simulations and in our segmented trajectories. The resulting length distributions are well-matched with only the single parameter *θ* needing to be fitted (Fig. 2D), with the exception of small FLS lengths which is due to the initial start of FLS growth not being captured by our model. The model also reproduces the complex shape of the distribution of persistence time for the FLS growth/shrinkage phases (Fig. 2 supplement 1).

Our model for the growth velocity thus suggests that FLS transit between multiple redundant modes of growth and strictly interpreted suggests that two independent complexes consisting of two regulatory protein pairs (X_1_,X_2_ and X_3_,X_4_) can control the growth and shrinkage dynamics of actin structures. The data however indicates that there are at least three pairs of two regulators each, the strongest being N-WASP/Cdc42•GTP, TOCA-1/Diaph3 and Ena/VASP (Fig. 1D). We know biologically that there are other proteins such as the formin-like protein 3 (38) as well as the bundling protein fascin that are involved in filopodia assembly, plus the regulators of FLS disassembly that we have not measured. Reconciling with the complexity of the numbers of proteins and complexes that are present, we found that many combinations of regulatory factor complexes (N being the number of proteins in each and M being the number of complexes) give rise to a Laplace-shaped velocity distribution (as long as 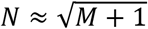 is roughly true, Fig. 2E, Fig. 2 supplement 1B,C). This argues that the actin bundle growth dynamics and shape distribution we observed can generically arise from redundant stochastic dynamics of actin regulators. The minimal model compatible with our data consists of a sum of three pairs each with a different fluctuation relaxation rate *θ* (the case of M = 2 and N = 3 in Fig. 2E, green box).

### Testing the dynamics of regulators and functional redundancy predicted by the theory

Using the example minimal case of *M* = 3 complexes of *N* = 2 regulatory factors, we can calculate that the relaxation rate *θ* scales as *λ*/*N*, with the overall velocity fluctuation relaxation rate *λ* ≈ 22 *min*^−1^ (determined from the fitted values of *θ* in Fig. 2 supplement 1). Hence, we expect *θ* to range around 11 min^−1^, which corresponds to a recovery half time of *τ* = ln(2)/*θ* ≈ 10 *s* (see supplementary information for further details). For simplicity in the calculations we have assumed that *θ* is the same for all proteins involved, in practice the dynamics will be dominated by the fastest component. To determine if the time scale *τ* from the fitting procedure agrees with the actual timescales of protein dynamics in FLS, we performed fluorescence recovery after photobleaching (FRAP) experiments (Fig. 3, Fig. 3 supplement 1). The recovery half times of the fastest components Diaph3 and VASP are 10-20 s, agreeing with theoretical predictions (actin and fascin are expected to recover quickly and completely as they move up from the membrane as the bundle grows, Fig. 3A-C). In addition, the same pairs of regulators that had the highest fluorescence intensity correlations with each other (Toca/Diaph3, N-WASP/GBD, Ena/VASP) also had a similar percentage recovery after photobleaching (Fig. 3A, C, Fig. 1D).

**Fig. 3.**
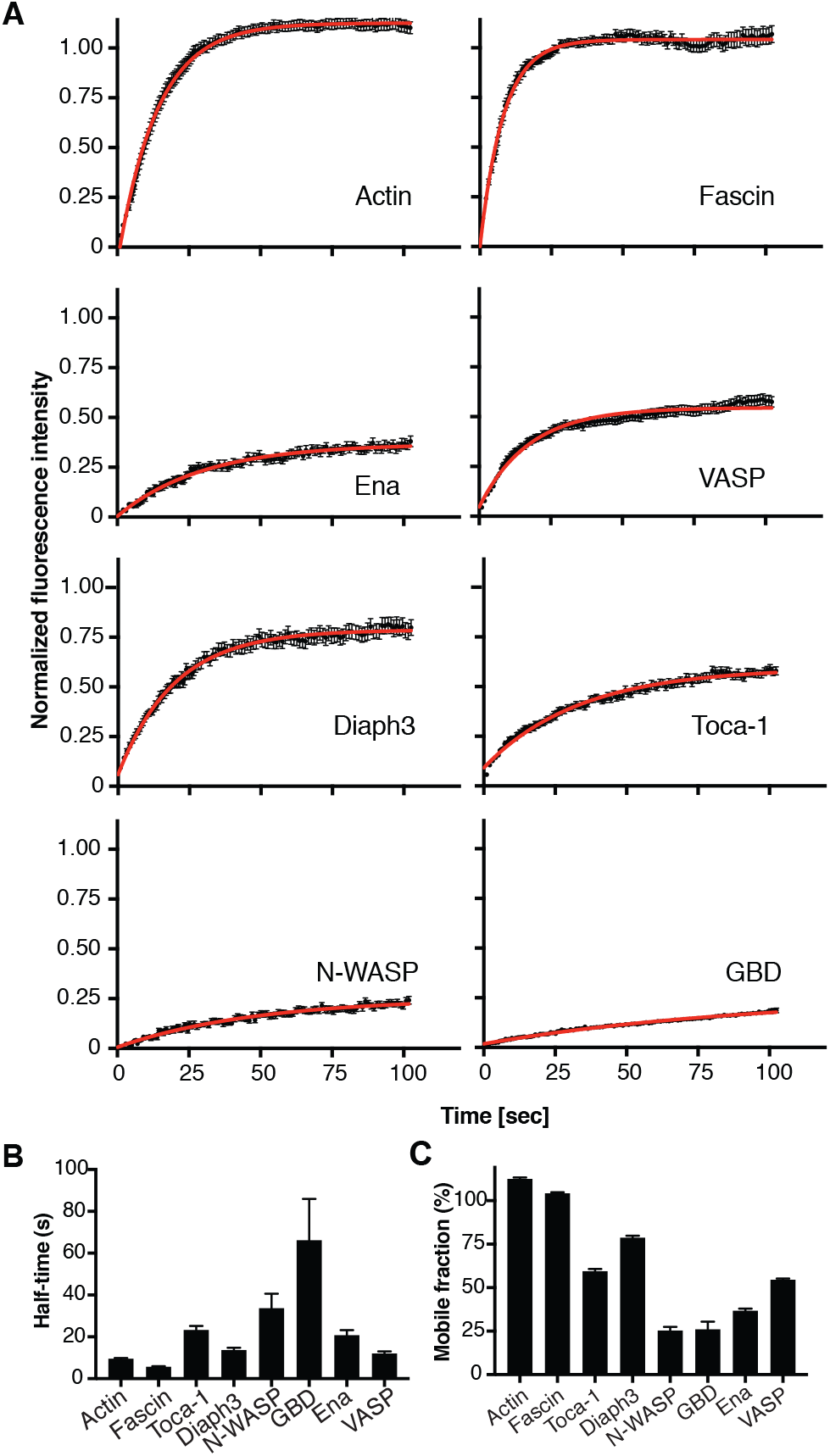
The FLS tip complex actin regulatory proteins is semi-dynamic. (A) Time course recovery of AF488 actin (n=55), GFP-Fascin (n=39), AF488 TOCA-1(n=65), GFP-Diaph3 (n=33), AF488 N-WASP (n=41), mKate-GBD (n=40), AF488 Ena (n=37), AF568 VASP (n=65) with fitted exponential (red) after photobleaching in tips of FLSs at steady-state grown for at least 30 minutes. Scale bar: 2 *μ*m. (B) The half-time and (C) percentage recovery from fitted exponential curves for each protein. Error bars represent standard error.

The model also implies that fluctuations of the growth velocity should correlate with the local concentration fluctuation of the regulators i.e. that there should be a change in fluorescence intensity of a regulator that precedes a change in filopodial velocity. To test this, we measured rapid timelapse stacks of the actin bundle growth and shrinkage at steady-state with the fluorescence intensity of actin regulators from within a pair of regulators (Ena and VASP) and between pairs (Ena/VASP to N-WASP). We tracked the correlation between changes in protein intensity level and growth velocity with time preceding extension of the FLS. There is a positive correlation (Fig. 4A), with a maximal correlation with a time offset of ~4 min prior to FLS extension, consistent with the need for actin incorporation to occur between the regulatory protein appearing and a change in FLS length (shaded areas are the 95% confidence interval). The negative correlations after t=0 show that a high change in velocity precedes a change in protein levels from the FLS tip. This is not a feature covered by the theory but would agree with actin regulators moving in and out of the tip complex.

**Fig. 4.**
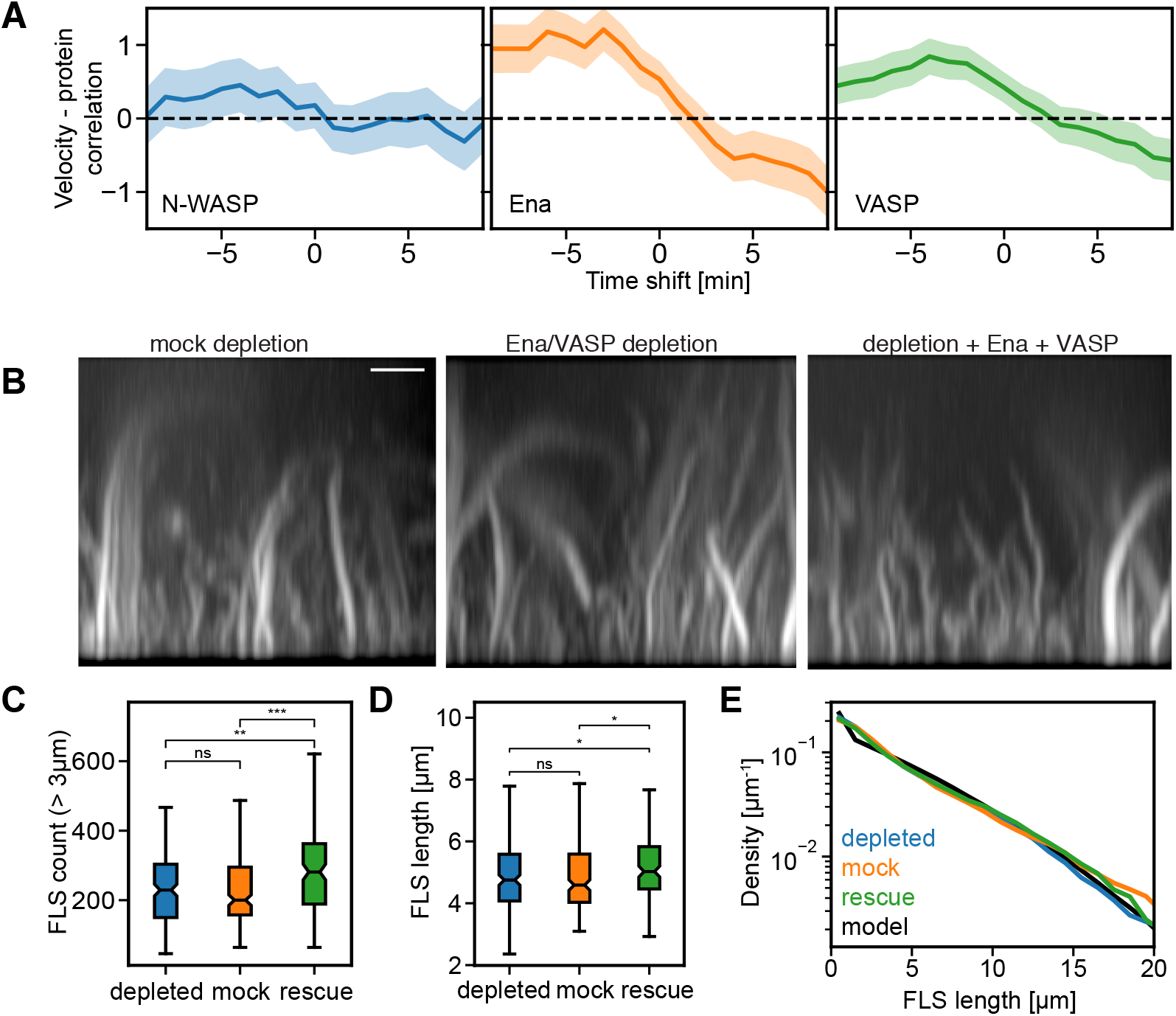
Velocity cross-correlations and effect of Ena/VASP immunodepletion predicted by the theory. (A) Time series cross-correlation of the absolute value of the FLS growth velocity with the absolute value of background-corrected N-WASP, Ena and VASP intensities at FLS tip complexes, averaged over all measured FLS trajectories. The graph shows the cross-correlation coefficient at the given time shift (negative time shifts mean that intensity changes precede velocity changes). Shaded areas indicate 95% confidence interval of the mean. (B) Side maximum intensity projections of confocal stacks of labeled actin in FLS assays with mock depleted extracts, Ena/VASP depleted extracts and depleted extracts with added 40 nM SNAP-Ena + 20 nM KCK-VASP (rescue). Scale bar = 5 *μ*m. (C) FLS numbers and (D) lengths with Ena/VASP depletion and rescue (E) Length distributions from Ena/VASP conditions, alongside calculated length distribution from reducing one subcomplex.

A more specific prediction of our model of stochastic and redundant FLS heterogeneity is that knocking out one of the subcomplexes should lead to almost negligible effects on the shape of the length distribution since this would only reduce the number of sum terms by one (Fig. 2E). To test this prediction experimentally, we co-immunodepleted Ena and VASP from our extracts. We observed no effect on the number or length of FLS, though adding additional Ena and VASP gave a very small yet significant increase in both length and number of FLS (Fig. 4B-D). The key feature in terms of the model is that the exponential distribution of FLS lengths is maintained (Fig. 4E). These observations are consistent with the data fitted by the numerical description of a shift from M = 3 to M = 2 independent pathways and with a redundant mechanism underlying FLS growth.

### Heterogeneous tip complexes and exponentially distributed filopodia lengths *in vivo* in *Drosophila*

Since FLS are an *in vitro* model we next tested whether the molecular characteristics FLS were similar to filopodia within a native context. Overexpression of proteins is expected to change the behaviour of actin dynamics that we are describing and we would also expect strong adhesion of filopodia to a substrate (such as in mammalian tissue culture) to alter their length distributions and dynamics relative to the native situation. For these reasons we turned to *Drosophila* because of the ability to knock-in fluorescent proteins into the endogenous locus and perform time-lapse microscopy within the physiological adhesive and mechanical properties of developing tissues where filopodia are naturally occurring. We used genome editing to knock-in a fluorescent protein into the endogenous locus of Ena *(enabled*, an actin filament elongation factor) and Arp2/3 complex nucleation promoting factor Scar/WAVE (*Scar*) and generated homozygous flies with both copies tagged (Ena is with GFP and Scar/WAVE with mNeonGreen). Using UAS GAL4 drivers to express the fluorescent membrane marker CD8- or CAAX-mCherry in specific cell types and 3D timelapse 2 colour laser scanning confocal imaging, we were able to visualize Ena and Scar/WAVE at the tips of filopodia in the leading-edge cells in dorsal closure (Fig. 5A,B) and in tracheal cells as the filopodia move (23, 39, 40) (Fig. 5C-D). The presence of Scar/WAVE at the filopodia tips supports a role of Arp2/3 complex activation in actin nucleation at these filopodia. For both proteins in both tissues we observed wide heterogeneity in protein localization. There are filopodia with a strong spot of Ena or Scar at filopodial tips, that tracks the filopodium (upper panels). There are instances where a spot is present but less bright or present for shorter times (middle panels), and there are instances where there is no spot present (lower panels). Both copies of the gene are tagged in the flies so there is no endogenous untagged protein present. The filopodia that have these different localization patterns can be adjacent to each other within the same cells (Videos 2-5). To quantify the data, we used maximum intensity z-projection timelapse stacks to visualize the filopodia through the z-stack and measured the maximum straight length observed in a filopodium’s lifetime. Through the timelapse series we looked for the brightest localization of Scar or Ena protein at the tip, in particular when the spot tracked the movement of the filopodial tip by cross-comparing between filopodial tips in the single confocal slices and z-projection. Once the appropriate slice was identified a region of interest was drawn at the tip corresponding to the diameter of the filopodium. Two background measurements were taken adjacent to the tip region of interest. The measurements confirm the wide heterogeneity seen by eye, and also that there is little correlation between the length of the filopodia and intensity of Ena or Scar fluorescence at the filopodia tip that are attained (Fig. 5 E, F). We applied our image analysis pipeline Filopodyan to determine the distributions of maximal filopodia path lengths, a more precise and reproducible measurement of length (41) and compared them with the analogous measurement of the distribution of FLS length trajectory maxima identified in our high time resolution timelapse movies. The filopodia lengths of both leading edge cells in dorsal closure and tracheal cells fell along exponential distributions together with *in vitro* FLS lengths (Fig. 5G), as we had observed above for FLS alone at steady-state (Fig. 1F) and also previously for myotubes in *Drosophila* (42).

**Fig. 5.**
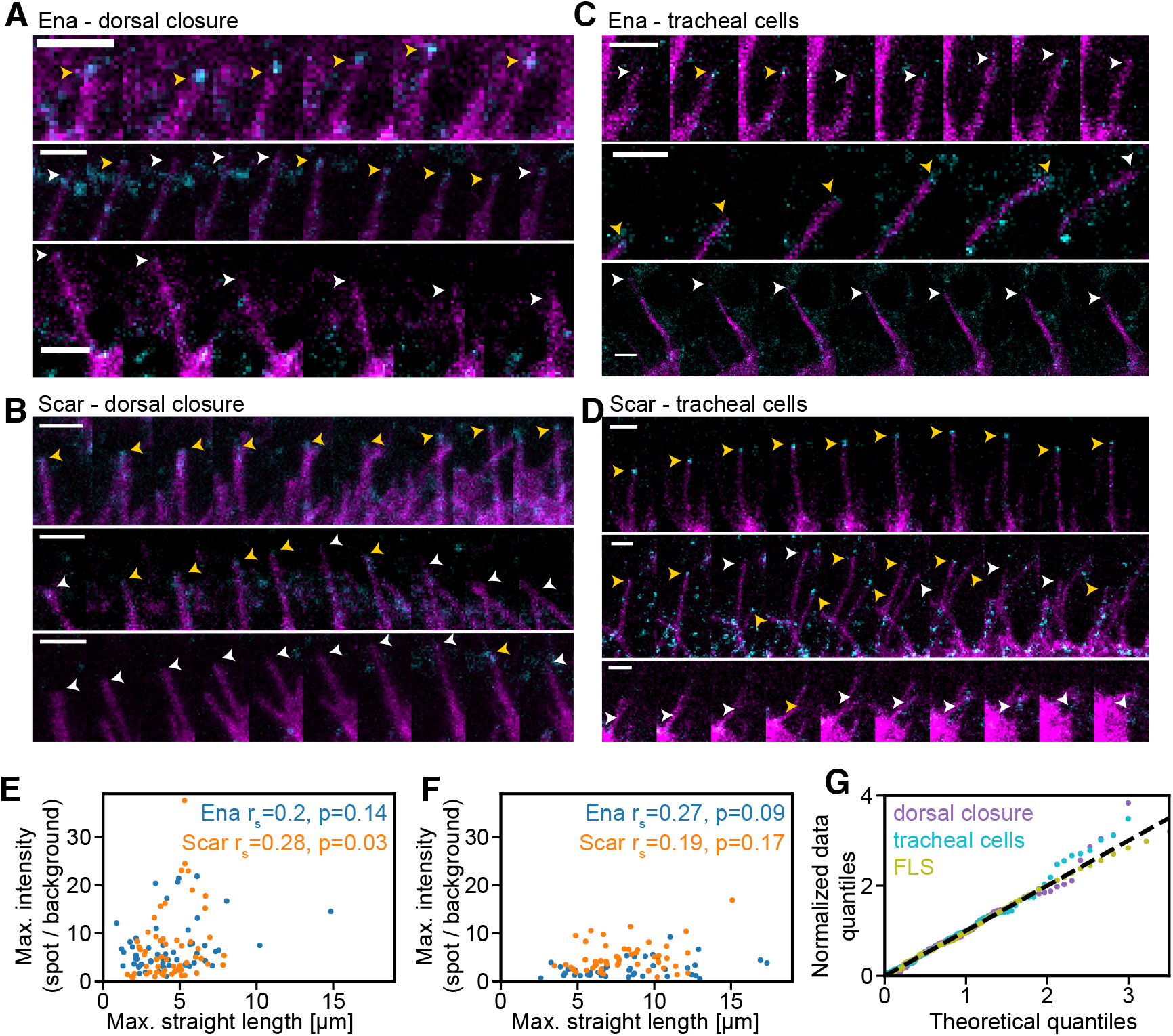
Heterogenous levels of actin regulators and exponentially distributed lengths of filopodia *in vivo*. (A) Timelapse montages of filopodia in leading edge cells in dorsal closure in the *Drosophila* embryo with a frame spacing of 15 seconds. The endogenous fluorescent protein Ena is shown in cyan, and magenta indicates the membrane marker expressed in the specific cell type using the Gal4 system. Pictures are maximum intensity projections of the cell marker together with the filopodial tip z-slice of the Ena channel. Yellow and white arrows indicate filopodia tips with and without protein, respectively. Scale bar = 2 *μ*m. (B) Same as (A) with endogenously labelled Scar. (C) Same as (A) for Ena in tracheal cells. (D) Same as (A) for Scar in tracheal cells. (E) Scatter plot of maximal Ena and Scar intensity at filopodia tips versus maximal end-to-end (straight) filopodia length in dorsal closure cells. Spearman correlation coefficients and associated p-values are given in the panel. (F) Same as (E) for tracheal cells. (G) Quantile-quantile plot showing distributions of filopodial lengths in *Drosophila* dorsal closure and tracheal cells as well as FLS lengths to the theoretical quantiles of an exponential distribution, the lines deviate when n is low. The lengths are taken at the time point at which a given filopodium or FLS is longest over its measurement lifetime.

In FLS we measured fascin at the tips (Fig. 1D), however fascin is also present in the shaft of both FLS and filopodia, and our theoretical framework does not rely on a tip localization. Even morphologically indistinguishable filopodia have vastly differing intensities of fascin in their shafts (Fig. 6A). We measured maximal fascin intensities through the shaft of FLS and filopodia in dorsal closure (using Filopodyan), finding a distribution of intensities similar in shape between filopodia and FLS (Fig. 6 B,C). This indicates that the function of fascin is similar in both, and shows that the FLS system captures genuine features of filopodia. We also observe fascin coming and going in the shaft of filopodia, originating from the tip, in agreement with fluctuations in its contributions (Videos 6 and 7). Again, we see no correlation between the maximal lengths of filopodia and their fascin intensity (Fig. 6D). To quantify the effects of perturbing fascin in filopodia, we measured the length distributions of filopodia during *Drosophila* dorsal closure under conditions of loss of fascin using the *singed* mutant or gain of fascin by overexpression of GFP-fascin using the engrailed-GAL4 driver. In *Drosophila* there is little effect on the mean lengths of filopodia from reducing the levels of fascin, where the loss of both fascin and the alternative bundling protein forked had a greater effect (40). Our model predicts that fascin null filopodia should be exponentially distributed but that overexpression should lead to a deviation in the length distributions from exponential as this would make the effect of a single complex dominant, equivalent to setting M = 1. The length distribution of dorsal closure filopodia from control embryos, GFP-fascin knock-in filopodia as well as filopodia in the fascin mutant embryos were well-fitted by exponentials (Fig. 6E,F). The overexpression data deviates markedly from an exponential distribution, in contrast to all cases of filopodia length measurements we have examined so far (Fig. 6E,F).

**Fig. 6.**
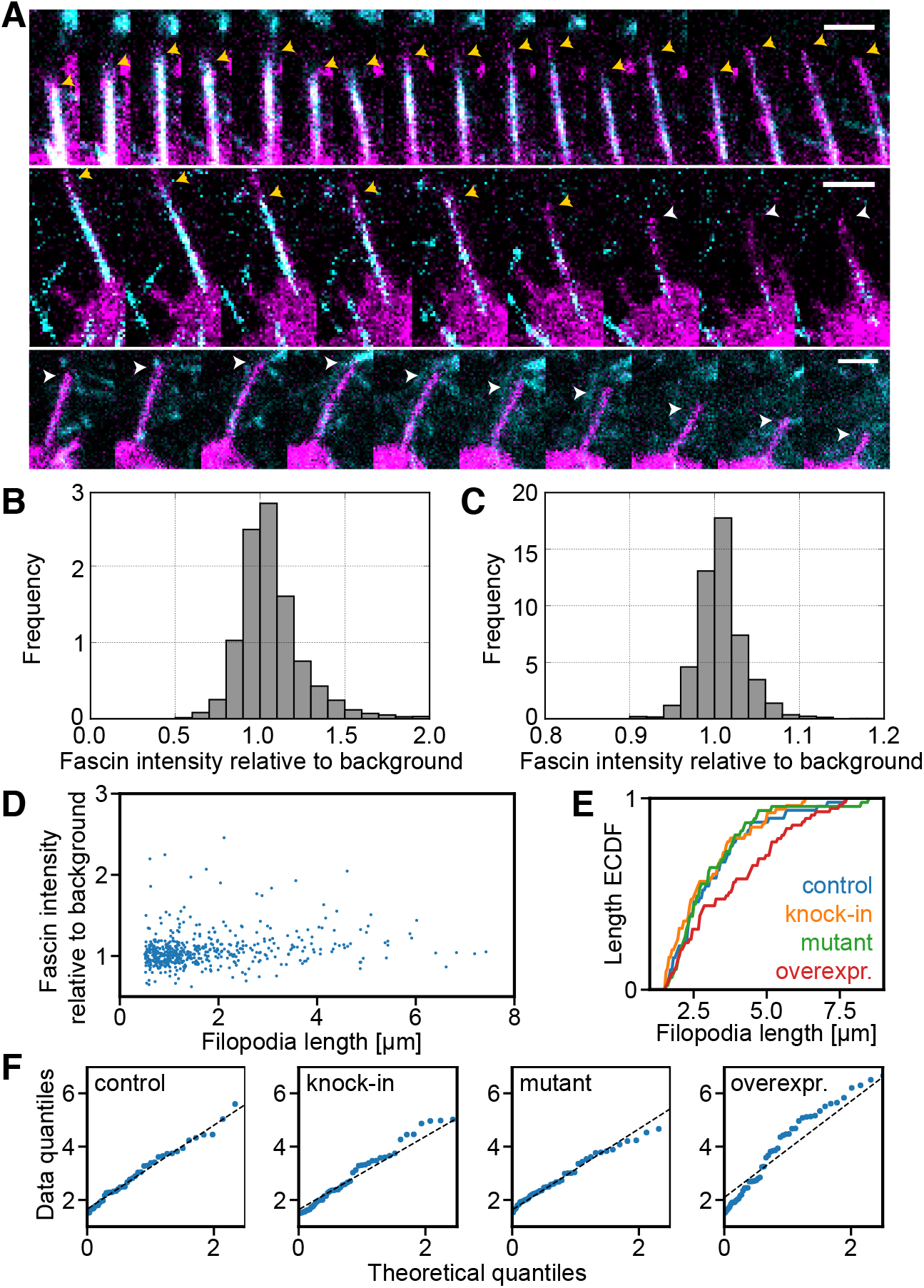
Heterogenous levels of actin regulators and exponentially distributed lengths of filopodia *in vivo*. (A) Timelapse montages of filopodia in dorsal closure cells in the *Drosophila* embryo with a frame spacing of 15 seconds. The endogenous fluorescent protein fascin is shown in cyan, and membrane marker in magenta. Pictures are maximum intensity projections. Yellow and white arrows indicate filopodia with and without fascin, respectively. Scale bar = 2 *μ*m. (B) Histogram of fascin intensity relative to local background in filopodia of dorsal closure cells. (C) Same as (B) for fascin intensity relative to local background in FLS shafts. (D) Scatter plot of fascin intensity in filopodia shafts versus maximal filopodia length in leading edge cells in dorsal closure. (E) Cumulative frequency plot of filopodia lengths with mutant, wildtype, knock-in GFP-fascin and overexpressed GFP-fascin in dorsal closure leading edge filopodia. p-values compared to control: labelled fascin p=0.765, mutant fascin p=0.996, fascin overexpression p=0.047 (Kolmogorov-Smirnov test). (F) Q:Q plots of filopodia lengths show control, labelled (knock-in) fascin and mutant fascin filopodia lengths are consistent with an exponential distribution but not fascin overexpression.

## Discussion

We have found that actin regulatory proteins form heterogeneous semi-dynamic assemblies on membranes comprised of at least 3-4 different subcomplexes where actin bundles nucleate and grow. At endogenous levels any individual protein has a limited capacity to dictate filopodial properties, with high redundancy, which gives rise to a robustness in overall actin dynamic properties. This explains our initially puzzling findings of weak, but positive, correlations between all actin regulators and bundle lengths, as well as the generic occurrence of exponential distributions of FLS lengths, independent of exact molecular composition. The subcomplexes we identified are reminiscent of proteins and interactions that are previously thought to be important in filopodia formation. Cdc42•GTP was most highly correlated with VASP and N-WASP (17, 43), Ena and VASP correlated with each other (33), and Diaph3, previously implicated in *de novo* filopodia nucleation, correlated with membrane-adaptor protein TOCA-1 (34).

The correlated positive fluctuations we observed between Ena, VASP and N-WASP intensity and FLS growth, and also the heterogeneity, are similar to those we have observed in growth cone filopodia (41). The heterogeneity we observe here is also similar to observations made regarding actin regulators that are used during clathrin-mediated endocytosis in mammalian cells (44). While it is thought that membrane tension is a determinant of whether the actin cytoskeleton is co-opted to help deform in the membrane during endocytosis, there is also variability between different regulators (44), suggesting a similar redundant mechanism may also operate during clathrin-mediated endocytosis.

Semi-dynamic complexes held together by networks of multivalent weak interactions are a fundamental principle of cellular organization (28). Recent studies of diverse cell biological structures, including P-granules, transcriptional complexes and actin signaling networks, have shown how these exist in semi-dynamic states phase-separated from the rest of the cytosol. Typically held together by networks of weak multivalent interactions between proteins and/or other molecules like RNA or membranes, they can also transition into glassy solids or amyloids. In FLS, the membrane interactions together with SH3 domain and proline-rich regions in Ena, N-WASP, VASP and Diaph3 are likely to contribute to the molecular interactions, similar to observations with N-WASP and Nck in purified systems (25, 26). While work in fibroblasts has indicated that the filopodia tip complex could be thought of as immobile (10) in neuronal filopodia VASP is partially mobile, similar to observations for FLS (45).

The redundancy in molecular composition described here may allow a variety of upstream and downstream components to intersect with control of actin, to bring the power of actin remodelling to different biological needs, while conserving the overall properties of the cytoskeletal behaviour. A multi-component system could also ensure that signals regulating the cytoskeleton must be multiple and coincident, as only rarely will a single input be sufficient to cause an effect, and it takes an overexpression scenario to subvert the normal homeostatic mechanisms, such as fascin in cancer (4). We show here that in spite of a dynamic and heterogeneous tip complex, it is possible for a constraint to emerge in the resulting activity such that actin cytoskeletal dynamics are similarly governed under a wide range of molecular states. These properties may be what allow the powerful actin machinery to be co-opted wherever necessary without altering its underlying properties.

## Materials and Methods

### Plasmids

*X. tropicalis* TOCA-1 (accession number BC080954) was PCR amplified from IMAGE clone 6980375 (Source Bioscience), and cloned into pET SNAP precision FA. *X. tropicalis* N-WASP (WASL) (accession number BC067309) was PCR amplified from IMAGE clone 5379332 (Source Bioscience), and cloned into pCS2-his-SNAP-FA. ZZ-TEV-WIP (Ho et al., 2004) was a kind gift from the Kirschner lab (Harvard, USA). *X. laevis* VASP (accession number BC077932) was PCR amplified from IMAGE clone 5515353 and cloned into pET SNAP precision FA with additional residues KCK added before the ATG, and site directed mutagenesis at GA192/193 to destroy endogenous cysteine. *X. laevis* Ena (accession number BC073107) was PCR amplified from IMAGE clone 5379332 and cloned into pCS2 his SNAP acceptor. *X. laevis* fascin (accession number BC097600) was PCR amplified from IMAGE clone 4970584 and cloned into pCS2 his GFP (GFP was PCR amplified flanked with ecoRI/fseI restriction sites and cloned into pCS2 his FA acceptor). Human N-WASP GBD domain was PCR amplified from pCS2-mRFP-GBD (a kind gift from William Bement, Addgene plasmid 26733) with XhoI/AscI flanking restriction sites and cloned into pET-pmKate2 to generate mKate-GBD. *X. laevis* Diaph3 was synthesised by Lifetechnology based on sequence alignments made with Mayball cDNA information and cloned into pCS2 his GFP (GFP was PCR amplified flanked with ecoRI/fseI restriction sites and cloned into pCS2 his FA acceptor).

### Protein purifications

Unless otherwise indicated, chemicals were purchased from Sigma-Aldrich, all steps were performed at 4 °C, purified proteins were concentrated using a 10,000 MWCO spin concentrator (Millipore), their concentrations were determined based on their absorption at 280 nm, and they were stored at −80 °C in 10 % glycerol following snap freezing in liquid nitrogen. To purify 6His-SNAP-TOCA-1, 6His-mKate-GBD, 6His-SNAP-Ena, 6His-GFP-fascin, 6His-GFP-utrophin-CH, 6His-GFP-Diaph3, pCS2 constructs were transfected into 293F cells by 293fectin reagent (Thermo Fisher Scientific) according to manufacturer’s instructions 48 hours prior to harvesting. pET and pGEX plasmids were transformed into BL21 pLysS *E.coli* and induced overnight at 19°C. Cells were harvested and pellets were resuspended in buffer containing 150 mM NaCl, 20 mM HEPES pH 7.4 with 2 mM 2-mercaptoethanol and EDTA-free cOmplete protease inhibitor tablets (Roche) prior to lysis by probe sonication. Following ultracentrifugation (40,000 rpm for 45 mins in a 70Ti rotor), proteins were affinity purified on Ni-NTA agarose beads (Qiagen). Proteins eluted from Ni-NTA beads by stepwise addition of increasing concentrations of 50-300 mM imidazole in a buffer containing 20 mM HEPES pH 7.4, 150 mM NaCl and 2 mM 2-mercaptoethanol. For all proteins apart from Diaph3, fractions were pooled and purified further using S200 gel filtration on an AKTA FPLC (GE Healthcare) in a buffer containing 150 mM NaCl, 20 mM HEPES pH 7.4 and 5 mM DTT. Diaph3 was used directly after affinity purification without further steps to prevent loss of activity.

To purify 6His-KCK-VASP, pET15b vector containing the 6His-KCK-VASP insert was transfected into Rosetta DE3 pLysS cells and induced overnight at 19°C. Cells were harvested and pellets were resuspended in buffer containing 300 mM NaCl, 20 mM HEPES pH 7.4 with 2 mM 2-mercaptoethanol and EDTA-free cOmplete protease inhibitor tablets (Roche) prior to lysis by probe sonication. Following ultracentrifugation (40,000 rpm for 45 mins in a 70Ti rotor), proteins were affinity purified on cobalt agarose beads (Talon superflow, GE Healthcare). Proteins eluted from cobalt beads by stepwise addition of increasing concentrations of 50-300 mM imidazole in a buffer containing 20 mM HEPES pH 7.4, 300 mM NaCl and 2 mM 2-mercaptoethanol. Fractions were pooled and concentrated using a 10,000 MWCO spin concentrator (Millipore). To purify 6His:SNAP:N-WASP/ZZ:WIP, the complex was expressed in 293F cells as described. The ZZ-WIP contains a TEV cleavage site between the tag and the protein. Lysates were incubated with IgG sepharose 6 beads (GE Healthcare) in XB buffer (100 mM KCl, 0.1 mM CaCl2, 1 mM MgCl2, 10 mM HEPES pH 7.4). Following extensive washing, TEV protease was added in XB buffer containing 10 mM DTT and 10 % glycerol and incubated overnight to cleave the N-WASP-WIP complex from the beads. The disposable column was drained to collect the cleaved protein complex, and applied to a small volume of GS beads to sequester the TEV protease. For SDS-PAGE analysis, samples were boiled in SDS sample buffer, run on 4-20 % polyacrylamide gels (BioRad) according to manufacturer’s instructions and analyzed by adding InstantBlue (Expedeon).

### Chemical labelling of proteins with fluorescent dyes

Labelling of SNAP-tagged recombinantly expressed proteins (6His-SNAP-Toca, 6His-mKate-GBD, 6His-SNAP-Ena, 6His-SNAP-N-WASP/ZZ-WIP) were performed with 5-10 *μ*M final protein concentration and 10 *μ*M dye (SNAP-Surface Alexa Fluor488 (New England Biolabs S9129S) or SNAP-Surface Alexa Fluor647 (New England Biolabs S9136S), respectively) in 50 *μ*l final volume with a buffer containing 150 mM NaCl, 20 mM HEPES (pH 7.4), 1 mM DTT and 1% TWEEN 20. After incubation overnight at 4 °C with protection from light and gentle rotation, excess dye was removed by two rounds of dialysis over 24 hours using a 0.1 ml Side-A-Lyzer MINI Dialysis Device with a molecular cut off of 20 kDa (10237043, Thermo Scientific), in a buffer containing 150 mM NaCl, 20 mM HEPES (pH 7.4) and 10% glycerol. Labelling of recombinantly expressed 6His-KCK-VASP was performed with approximately 50 *μ*M final protein concentration and 10 – 20-fold molar excess of maleimide dye (AF 568 C5 Maleimide (Life Technologies, A-20341) or AF 647 C5 maleimide (Life Technologies A-0347), respectively) in a 10-fold molar excess of TCEP. After incubation overnight at 4 °C with protection from light and gentle rotation, the dye was removed by buffer exchange using a spin concentrator (Amicon Ultra-15 Centrifugal Filter Unit with an Ultracel-10 membrane (Merck Millipore, UFC901024), and a buffer containing 20 mM HEPES (pH 7.4), 300 mM NaCl and 10% glycerol. Unlabelled skeletal rabbit muscle actin was prepared as described (Daste et al., 2017).

### Quantitative Western blotting

A dilution series of *X. laevis* egg extracts, as well as a dilution series of recombinant purified proteins were size-separated by SDS-PAGE. Samples were transferred to a nitrocellulose membrane by wet transfer using a Bio-Rad Mini Trans-Blot Cell apparatus using transfer buffer (25 mM Tris, 192 mM Glycine, 0.1% SDS, 20% methanol) for 1h at 0.38 A. Blots were rinsed with deionised water and blocked in TBST with 5% milk (skimmed, household milk powder) for 20 min at room temperature. Primary antibodies were diluted in 5 ml blocking buffer each and incubated with the membrane for 1 h at room temperature or overnight at 4 °C. Blots were imaged on a LICOR BioSciences Odyssey Sa scanner. Intensities of the dilution series of the recombinant protein with known concentrations were measured and used as standard curve for the extract samples. Blots were repeated at least twice for each protein. The Toca-1 antibody (Ho et al., Cell 2004), VASP antibody and fascin antibody (Lee et al., 2010) were a kind gift from Marc Kirschner, Harvard, USA. The N-WASP antibody was raised against full length purified *X. tropicalis* 6His-SNAP-N-WASP/ human ZZ-WIP in rabbit and affinity purified against the same protein sample. The Ena antibody was raised against full length purified *X. laevis* 6His-SNAP-Ena in rabbit and affinity purified against the same protein sample.

### FLS assays

Supported lipid bilayers and high speed supernatant frog egg extracts were made and FLS assays performed as previously described (Walrant et al., 2015). For snapshot and time-lapse data, a 1:6 dilution of 25 mg/ml *X. laevis* egg extracts was used, as optimized for FLS spacing and image segmentation by FLS Ace. For FRAP experiments, a 1:4 dilution of *X. laevis* egg extracts was used, as optimized for ROI selection on the membrane. Typically, a final volume of 50 *μ*l of reaction mix (2 mM DTT, 1 x energy mix (50 mM phosphocreatine, 20 mM Mg-Adenosine triphosphate (MgATP), 1 x XB buffer (100 mM KCl, 100 nM CaCl_2_, 1 mM MgCl_2_, 10 mM K-HEPES (pH 7.4), 50 mM sucrose), 4.2 mg/ml HSS *X. laevis* egg extracts, labelled proteins of interest and 10 x XB to account for salt concentration), stored on ice, was gently added to the supported lipid bilayer at room temperature. We optimised our microscopy to add as little recombinant protein as possible while giving acceptable signal/noise in the images to minimize the influence of the added labelled protein. For measurements of protein intensities at tip complex areas, recombinantly expressed and chemically labelled proteins were added in the following final concentrations: Actin (labelled) 210 nM; TOCA-1 10 nM; VASP 20 nM; N-WASP 20 nM; Fascin 300 nM; GBD 2 nM; Diaph3: 20 nM, Ena: 40 nM. For FRAP experiments, recombinantly expressed and chemically labelled proteins were added in the following final concentrations: Actin (labelled) 210 nM; TOCA-1 48 nM; VASP 175 nM; N-WASP 18 nM; Fascin 650 nM; GBD 2.46 nM; Diaph3 16 nM; Ena 60 nM. For snapshot images, a final concentration of 10 *μ*M of unlabelled rabbit skeletal muscle actin was added to the reaction mix. For time-lapse imaging, unless stated otherwise, a final concentration of 1 *μ*M of unlabelled rabbit skeletal muscle actin was added to the reaction mix. For velocity measurements, actin dynamics were monitored by addition of 250 nM GFP-utrophin-CH domain instead of labelled actin to prevent bleaching. For time-lapse videos, microscopic measurements were started 3 min post initiation. For snapshot images microscopic measurements were performed at 20-30 min, and for FRAP experiments 30-40 min post initiation.

### Immunodepleted extracts

For immunodepletion of endogeneous Ena and VASP from *X. laevis* egg extracts, three batches of protein A dynabeads (Life Technologies, 100-02D) were coupled to respective custom raised and affinity purified antibodies in PBST (PBS + 0.02 % TWEEN 20) for 1 h at room temperature under end-over-end rotation. Beads were washed three times with 500 μl PBST, then three times with 500 μl XB. *X. laevis* egg extracts were subjected to three round of incubation with coupled beads for 30 minutes with end-over-end rotation at 4 °C. For Ena and VASP double depletions, Ena and VASP antibodies were coupled to beads in separate reactions and combined prior to adding extracts. A rabbit IgG polyclonal isotype control antibody (Abcam, ab27478) was used for mock depletions.

### Fluorescence microscopy

Microscopic images were acquired on a custom combined total internal reflection fluorescence (TIRF)/spinning disc system supplied by Cairn Research based on a Nikon Eclipse Ti-E inverted microscope equipped with an iLas2 illuminator (Roper Scientific), CREST X-light Nipkow spinning disc, 250 μm NanoScanZ piezo driven Z-stage/controller and Lumencor Spectra X LED illumination using a 100x 1.49 NA oil immersion objective. Images were collected at room temperature with a Photometrics Evolve Delta EM-CCD camera in 16-bit depth using Metamorph software Version 7.8.2.0 (Molecular Devices). Atto-390, AF 488, 568 and 647 samples were visualized using 460/50, 470/40, 560/25 and 628/40 excitation and 525/50, 585/50 and 700/75 emission filters, respectively. For fluorescence recovery after photobleaching experiments, FLS were imaged 30 min post initiation. A reference TIRF image of the protein of interest and a confocal Z stack of the actin channel were acquired. A maximum of eight circular ROIs of 20×20 pixels (2.97×2.97 *μ*m) (e.g. four FLS tips and four corresponding background regions) were selected. Five frames were recorded in 1 second intervals prior to bleaching using the 488 nm or 561 nm laser at 25% power for two iterations. Fluorescence recovery was recorded by the corresponding laser line at 5% laser power for 120 frames, with intervals of 0.5 sec for the first 40 frames and 1 sec for all following frames. Changes in fluorescence intensity in the ROI after photobleaching were analyzed in ImageJ using a publicly available script developed at the Image Processing School Pilsen 2009 according procedures previously described (46). The bleach frame was identified by the largest frame-to-frame drop in intensity. For each analysis, four ROIs; bleached FLS (FFLS), non-bleached FLS (nFFLS), bleached background (FBG), and non-bleached background (nFBG) were specified. The non-bleached regions were selected in close proximity to the corresponding bleached regions, to account for local background. Firstly a background ROI intensity subtraction from the measured values was performed on the raw data, and the pre-bleach frames were normalized to 1. Loss of fluorescence during post-bleach acquisition was corrected for based on the unbleached ROI, and furthermore the bleach frame was normalized to 0.

### FLS Image Analysis

For extracting data from spinning disc confocal and HILO microscopy images of FLS, we developed FLS Ace, a plugin for Fiji (47) that maps actin structures in z-stacks and measures signal intensity in images representing other channels. FLS are segmented using a 2D Difference of Gaussians filter to define positions independently in each XY plane which are then traced through Z starting from each position found in the base plane using a greedy algorithm. The plugin has a user-friendly interface allowing the base plane, DoG sigma and k values, thresholding method, minimum required length for traced FLSs and the maximum tracing radius to be set and tested on a single actin stack to ensure accurate FLS detection. These parameters can then be used in batch mode to map FLSs in many actin stacks taken at a series of timepoints and measure as many associated images as required. Linear assignment between structures detected in multiple timepoints allows for phenotypic analysis of individual FLS throughout their lifetime. Batch mode is parallelized for speed and an XML configuration file is used to define gene names, regular expressions to determine which images correspond to each gene and the segmentation parameters to use for each. The actin-based segmentation is used to measure signal intensity in FLS bases in any additional (TIRF) channel. To alleviate systematic spatial shifts stemming e.g. from chromatic aberration, we performed translation image registration using the actin base slice as a reference. Local background in each channel is determined by averaging the fluorescence intensity in a ring two to three times the size of the FLS base, excluding any neighboring FLS bases. In order to efficiently process our GFP-utrophin-CH-based high time resolution time lapse Videos, we re-implemented the above algorithm in Python using algorithms from scikit-image.

### Post processing of snapshot data

We filter out segmented FLS with an effective diameter of less than 0.5 *μ*m because base size information as well as fluorescent intensity measurements become unreliable at or below the resolution limit. When considering FLS length distributions, we truncated the distribution to lengths between 5 *μ*m and 20 *μ*m to exclude structures which have not yet matured into FLS and artefacts stemming from the lower z-resolution in our confocal stacks. Fluorescence intensities of observed tagged proteins are determined by subtracting the local background intensity from the intensity averaged over all pixels in the FLS base area. To determine protein-protein and protein-morphology correlations, we calculate the Spearman correlation values for each field of view separately and average over the resulting ensemble.

### Post processing of time lapse data

Artificial breaks in trajectories can occur due to segmentation errors. We took care to repair these breaks by merging trajectories with average base positions closer than 1 *μ*m, within 6 time points and no temporal overlap, using a greedy algorithm. To reduce noise from image segmentation errors, we smoothed FLS length and growth velocity trajectories using a Savitzky-Golay filter of order 3 with a window of 11 time points.

### *Drosophila* stocks

Flies were raised and crossed at room temperature. The wild-type strain used was *white[1118]*. Fascin mutant *sn[28]* and the transgene *UAS-GFP-Fascin* (kindly provided by Brian Stramer) are described in FlyBase. *Mef2-Gal4* was used to drive in myotubes, *Btl-Gal4* in trachea and *engrailed-Gal4* in the leading edge cells in dorsal closure. To make *GFP-fascin; en-Gal4 UAS-cd8mCherry / (CyO)* flies, the original flies used for knock-in were *y[1]sc[1] v[1];;{y[+t7.7] v[+t1.8]=nanos-Cas9}attp2*. Fascin was tagged by CRISPR/Cas9 mediated genome editing with GFP at the N-terminus. GFP insertion was before the first amino acid of Fascin, with the addition of a linker sequence such that the fusion protein junction corresponds to TKA**SSSS**ATG (linker in bold). Two guide RNA sites were chosen, one at the GFP insertion point and the other 1.4kb upstream in the 5’UTR.

Donor plasmid: pTv-[w+] Fascin GFP constructed by In-Fusion cloning of PCR generated 5’ and 3’ homologous arms, GFP and ClaI/XhoI cut pTv-[w+] vector. 1.4kb 5’ homologous arm amplified from CFD2 genomic DNA with primers TATTCGAATCTGCAGCGTTTAGCGTTACTGACTGTGGGC and CTTTACTCATGGTGCTGATGGGAGCAATCT. 1.3 kb 3’ homologous arm amplified from CFD2 genomic DNA with primers TTCGAGTTCATCTATGAACGGCCAGGGCTGCGA and AATGGCACTGTTATCGCTATCATCTATTGAGCCATTTAGCCA. The gRNA sequence is split by the insertion of GFP so it was not necessary to introduce silent mutations to prevent cleavage of the donor plasmid. GFP was amplified with primers AGCACCATGAGTAAAGGAGAAGAAC and TTCATAGATGAACTCGAAGCTTTGTATAGTTCATC.

gRNA plasmid: Fascin gRNA target sequences were cloned into pCFD4-U6:1_U6:3tandemgRNAs plasmid (a gift from Simon Bullock (Addgene plasmid # 49411) by In-Fusion cloning (48). Primers TATATAGGAAAGATATCCGGGTGAACTTCGCTCCCATCAGCACCATGAAGTTTTAGAGCTAGAAATAGCAAG and ATTTTAACTTGCTATTTCTAGCTCTAAAACGCTCGCAGCCCTGGCCGTTCCGACGTTAAATTGAAAATAGGTC were used to amplify and introduce the two Fascin gRNAS from pCFD4-U6:1_U6:3tandemgRNAs which was then cloned into BbsI cut pCFD4-U6:1_U6:3tandemgRNAs.

CDF2 nos-Cas9 fly embryos (genotype: y1 P(nos-cas9, w+) M(3xP3-RFP.attP)ZH-2A w*, a gift from Simon Bullock) were injected with 250 ng/ul each of donor and gRNA constructs Male flies were crossed with C(1)DX balancer.

To make *Scar/WAVE-NeonGreen; en-Gal4 / CyO; UAS-cd8mCherry / TM2TM6* flies, SCAR/WAVE was tagged by CRISPR mediated genome editing with mNeon green at the C-terminal. mNeon green insertion was between the final amino acid of Scar/WAVE and the stop codon, with the addition of a linker sequence such that the fusion protein junction corresponds to TSASSSSATG (linker in bold). Two guide RNA sites were chosen which flanked the insertion site, with the PAM motifs being separated by 6 nucleotides.

Donor plasmid: pTv-[w+] Scar/WAVE mNeon green constructed by In-Fusion cloning of PCR generated 5’ and 3’ homologous arms, mNeon green and HindIII/XhoI cut pTv-[w+] vector. 1.2 kb 5’ homologous arm amplified from CFD2 genomic DNA with forward primer GAATCTGCAGCTCGATCAACGGCTCTAATATCTCACATTC and reverse primer CGATGAGCTCGAAGCTGAAGTTTCGTTCGGTTCCATCCAGCCCTCGC. The reverse primer has two silent changes from genomic sequence in the gRNA target sequence (C to T and T to A at positions 16 and 19) to prevent cleavage of the donor plasmid. 1.4 kb 3’ homologous arm amplified from CFD2 genomic DNA with forward primer CAAGTGATCCCTGATAAATTCGTTAAAGCCTG (with silent changes from genomic sequence of T to A and C to T at positions 16 and 19) and reverse primer TCGAAAGCCGAAGCTGCCCACCGCAATTAGCTTATATTG. mNeon green was amplified from pNCS mNeon green (Allele Biotech) with primers GCTTCGAGCTCATCGATGGTGAGCAAGGGCGAGGAGGATA ACATGGCCTC and ATCAGGGATCACTTGTACAGCTCGTCCATGCCCATC.

gRNA plasmid: Scar/WAVE gRNA target sequences were cloned into pCFD4-U6:1_U6:3tandemgRNAs plasmid by In-Fusion cloning. Primers TCCGGGTGAACTTCGGATGGAACCGAACGAAACATGTTTTAGAGCTAGAAATAGCA AG and TTCTAGCTCTAAAACTGATTAACTCGTTAAAGCCTCGACGTTAAATTGAAAATAGGTC were used to amplify and introduce the two Scar/WAVE gRNAS from pCFD4-U6:1_U6:3tandemgRNAs which was then cloned into BbsI cut pCFD4-U6:1_U6:3tandemgRNAs.

CDF2 nos-Cas9 fly embryos were injected with 250 ng/ul each of donor and gRNA constructs. Two independent lines were established with mNeon Green tagged Scar/WAVE: 15 X F0 male flies were crossed with Gla/CyO balancer (injected males were pre-screened by pcr with the forward 5’ homologous arm primer and a reverse mNeon green primer CACCATGTCAAAGTCC). 4 X Positive F1 flies (pre-screened as for F0 flies) were then crossed with Gla/Cyo balancer line. Correct integration of mNeon green was confirmed in two lines by PCR and sequencing across the entire donor sequence. PCR was also performed with primers flanking the donor sequence to confirm the size of the integrated fragment.

To make GFP-ena[w+]/CyO; UAS-cd8mCherry flies, Enabled was tagged with GFP in its endogenous gene locus using genomic engineering (49). After deletion of almost the entire Enabled ORF and 3’ UTR (2R: 19,158,510 – 19,163,715; release r6.07) including the N-terminal EVH1 domain by homologous recombination using a 5.3 kb 5’ and a 3.4 kb 3’ homology arm, it was replaced with an attP integration site (Ena knock-in platform: Ena[GX]). A wild-type Ena genomic rescue construct created by subcloning a 5,203-bp fragment from the Bac RP98-01N09 (Berkeley Drosophila Genome Project) was re-inserted, containing 224 bp upstream of the nucleotide corresponding to the first nucleotide of the Ena EVH1 domain, and 2025 bp downstream of the stop codon of Ena, into the attB vector pGE-attB-GMR (49). The GFP (variant mGFP6) coding sequence was inserted at the N-terminus of the EVH1 domain with a short linker amino acid sequence at the C-terminus of GFP, the fusion protein junction corresponds to YKASSSSEQS (linker in bold). GFP will be fused to all annotated isoforms of Enabled.

### Live imaging and data analysis of *Drosophila* embryo filopodia

As previously described (42) dechorionated embryos (washed in 50% bleach) were mounted on a glass-bottomed dish with heptane glue and submerged in water. To identify GFP-ena homozygous embryos balancer chromosome CyO with Dfd-YFP were used and we did negative selections for Dfd-YFP expression in the head region of embryos. Live microscopy was performed on an inverted Leica TCS-SP5 equipped with a 63x 1.4 NA Plan Apo oil immersion objective at room temperature. To visualize filopodial movement and respective protein intensities within filopodia in dorsal closure, embryos were imaged at the end of stage 14. Microscopic measurements were performed by taking z-stacks of 7-11 z sections (0.5 *μ*m spacing). Time-lapses were takes at 15 second intervals for 10 min. Ena and Scar/WAVE intensities at the tips of filopodia were quantified manually using ImageJ. Filopodia lengths and fascin intensities were quantified using our previously developed open-source pipeline Filopodyan (41). Using ImageJ, a maximum projection of the respective stacks was applied for filopodia reconstruction. A combination of automated detection with some manual editing was used to track identified filopodia over time.

## Acknowledgments

We would like to thank Bishara Marzook, Helen Fox and Louise Whiteley for protein preparation and immunodepletion assistance, Vivek Bhogadi and Sven Huelsmann in the creation of fascin and Ena GFP fly stocks and Nick Brown, Helen Mott, David Owen, Sandra Schmid and Marc Kirschner for critical reading of the manuscript. We thank the Gurdon Institute imaging facility for microscopy support. We dedicate this work to Guilherme Correia (Will) who started this project for his PhD and was tragically unable to continue.

## Author contributions

Conceptualization: JLG; Data curation: UD; Formal analysis: UD, EH, BDS; Funding acquisition: UD, JLG, BDS, EH, HS; Investigation: IKJ, HS, BR, YI, JM, JRG, AW, JLG; Methodology: IKJ, UD, EH, BR, YI; Project administration: JLG; Resources: YI, BR, JM, AW, JRG; Software: UD, IKJ, RB; Supervision: BDS, JLG; Validation: BR, JRG; Visualization: HS, IKJ, UD; Writing - original draft: IKJ, UD, JLG. Writing - review and editing: all authors.

## Data and materials availability

All data is available in the main text or the supplementary information. All data, code, and materials used in the analysis are available for the purposes of reproducing or extending the analysis. mNeon green constructs are supplied by the Allele Biotech repository.

---

**Fig. 1 - supplement 1.**
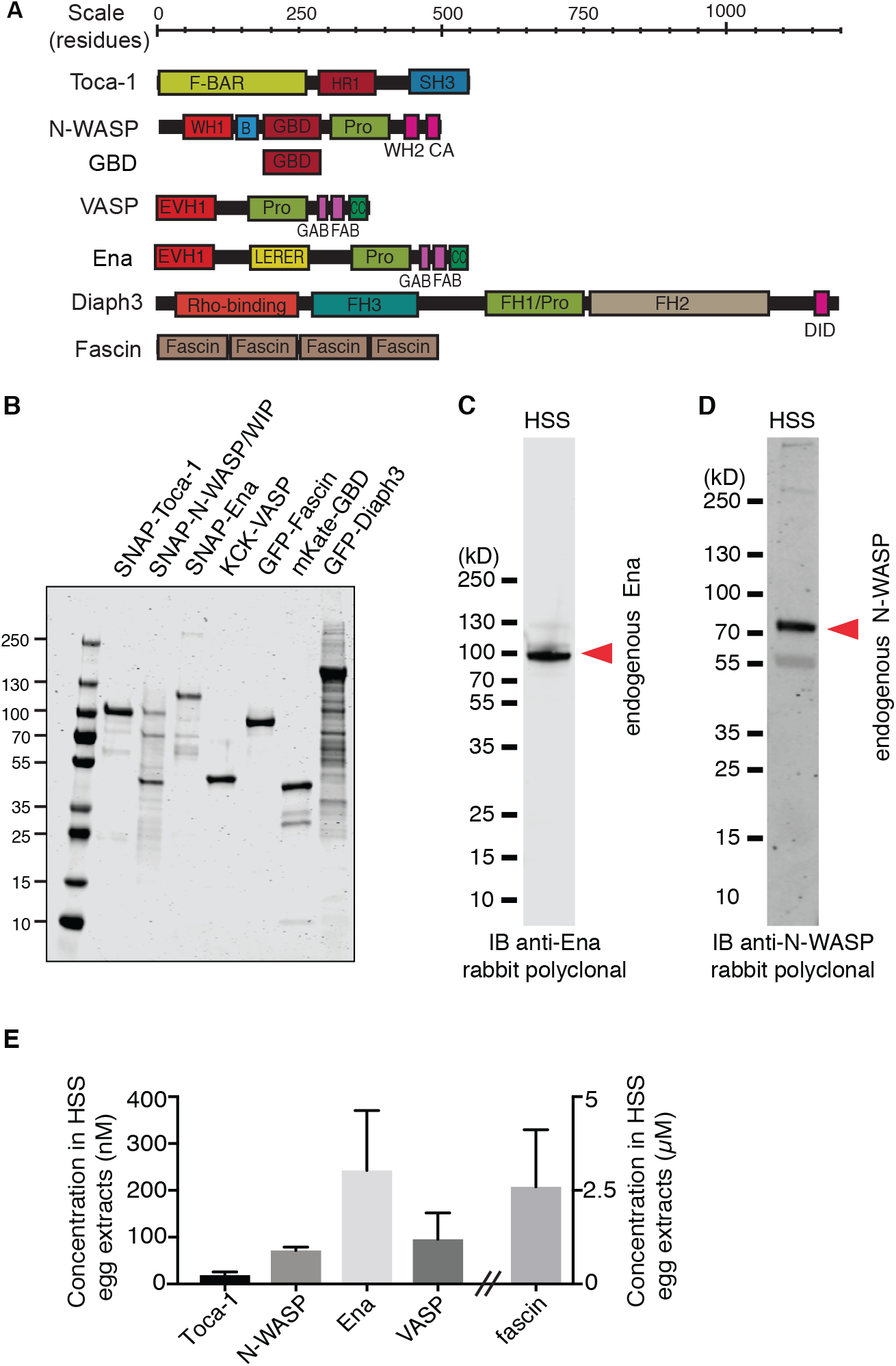
Purified protein gels and quantitative western blots. (A) Domain structures of actin regulatory proteins used in this work (B) Coomassie staining of purified proteins size separated via SDS-PAGE. (C) Western blot staining of extract sample size-separated via SDS-PAGE with anti-Ena rabbit polyclonal antibody raised against 6His-SNAP-ENA. (D) Western blot staining of extracts size-separated via SDS-PAGE with anti-N-WASP rabbit polyclonal antibody raised against 6His-SNAP-NWASP. (D) Quantification of protein concentration in *X. laevis* egg high speed supernatant extracts (HSS) by quantitative Western blotting, n = 3 (Toca-1, N-WASP) or n = 4 (Ena, VASP, fascin).

**Fig. 1 - supplement 2.**
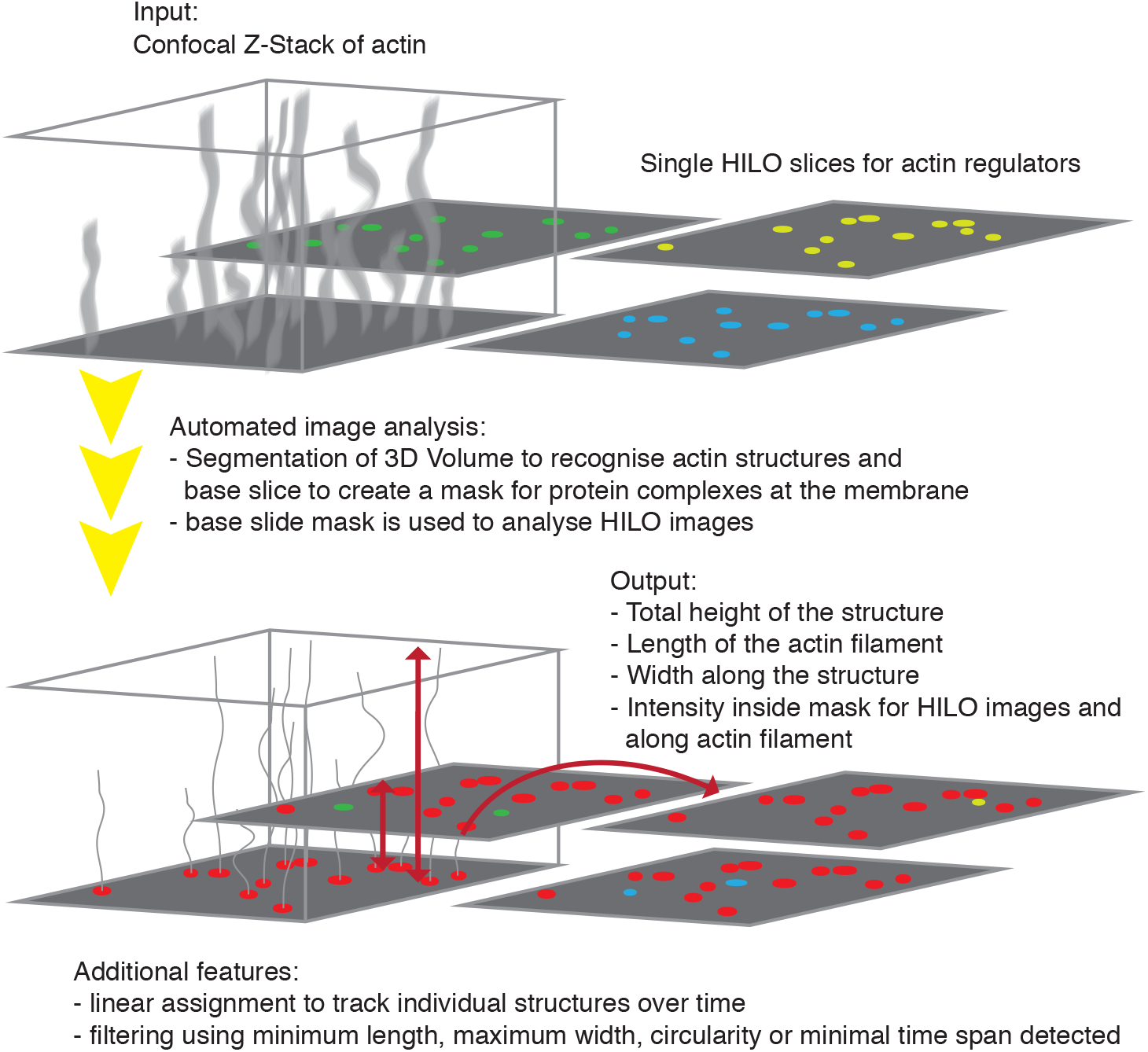
FLS Ace image analysis pipeline used to segment and measure FLS. Workflow of the custom image analysis pipeline implemented as a PlugIn for ImageJ. Microscopic images are obtained via HILO and confocal or widefield illumination (for z-stacks) on the same fields of view. Individual FLS are segmented based on the actin fluorescence from the z-stack. A mask created from this stack is used to overlay on any additional channel, measuring intensities inside the area defined as the actin regulatory complex assembly at the membrane with background correction (at the base slice), and along the shaft. Output includes protein intensity information as well as shape parameters.

**Fig. 1 - supplement 3.**
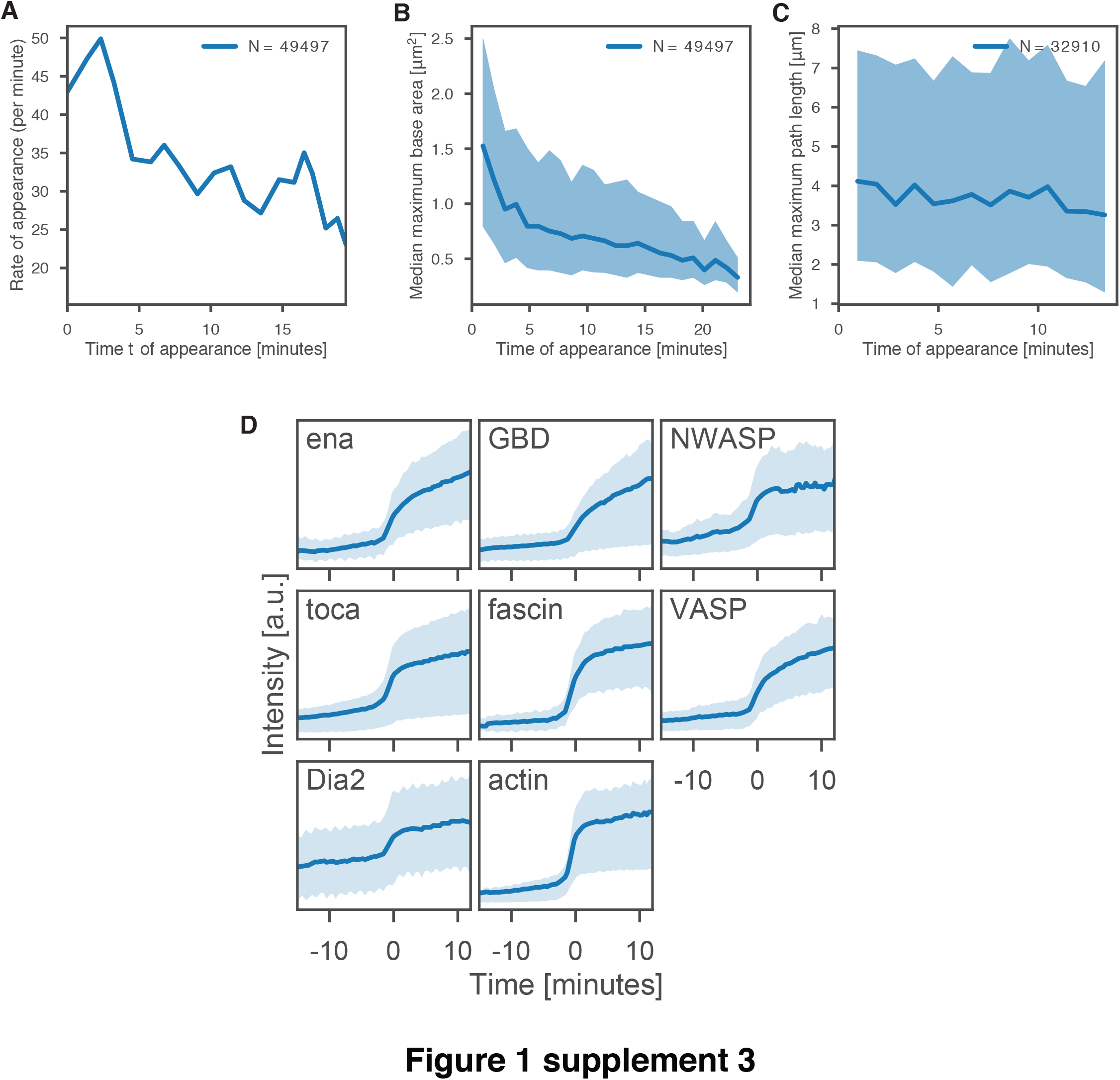
New FLS are created with an approximately uniform rate throughout the observation time. (A) Rate of appearance of FLS with time. (B) FLS that arise early in the experiment end up having a larger diameter (C) The lengths of FLS that arise early and late are similar. (D) Mean increase in protein intensity during FLS formation for each protein individually. Shaded areas in all graphs are the standard deviation.

**Fig. 1 - supplement 4.**
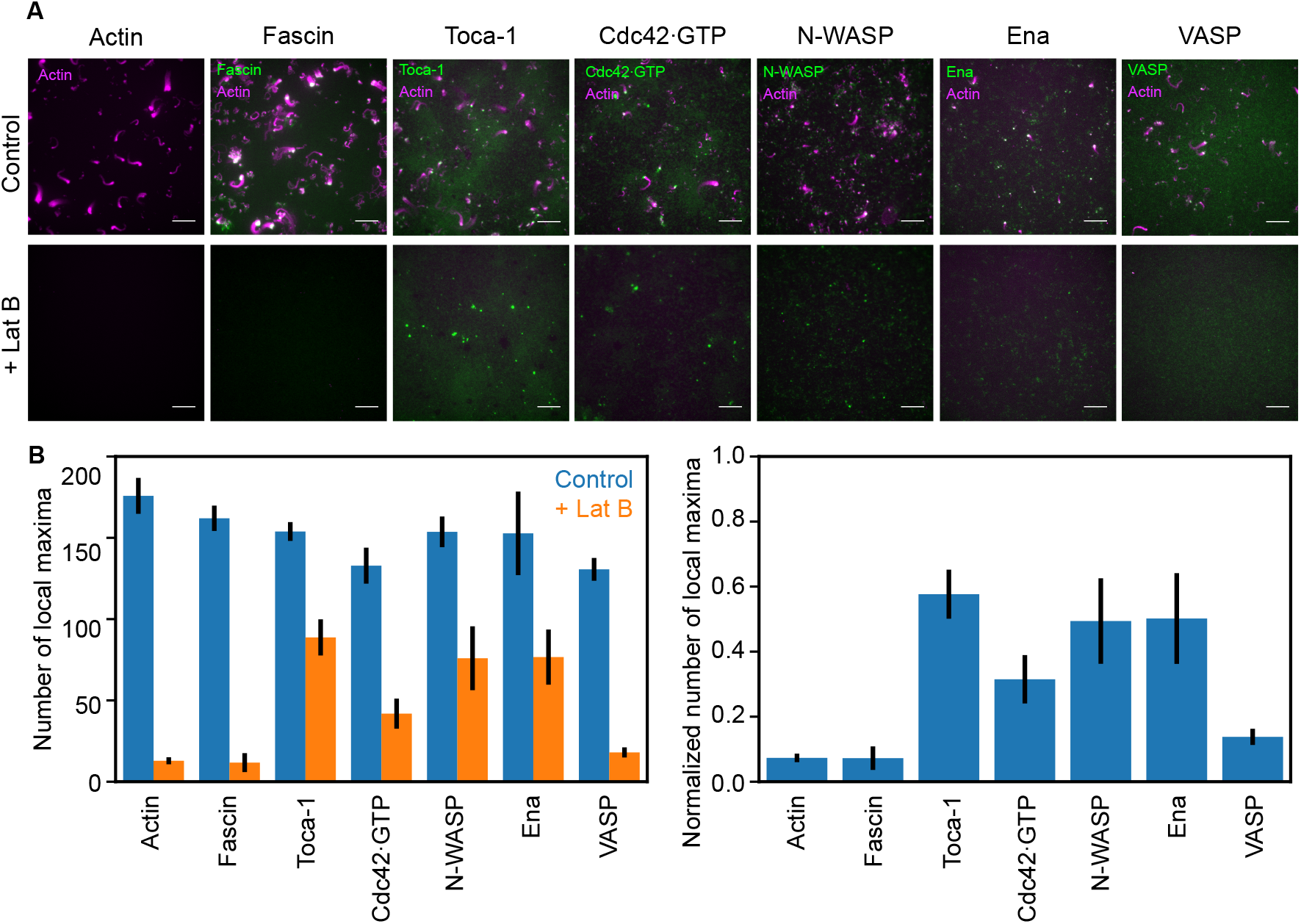
FLS tip complex assembly is primarily actin-dependent. (A) Latrunculin B treatment has some effect on the assembly of TOCA-1, Ena and N-WASP foci on the membrane, and leads to inhibited recruitment of actin, Fascin and VASP, with partial inhibition of Cdc42•GTP recruitment. Scale bar: 10 *μ*m. (B) Quantification of number of local maxima computed by protein fluorescence intensities plus and minus LatB. The right panel shows the fraction of local maxima when LatB was added normalized to the control. Data are from 3 or more independent experiments. Error bars indicate the standard error of the mean. All proteins were significantly reduced at the p < 0.05 level (Welch’s t-test).

**Fig. 1 - supplement 5.**
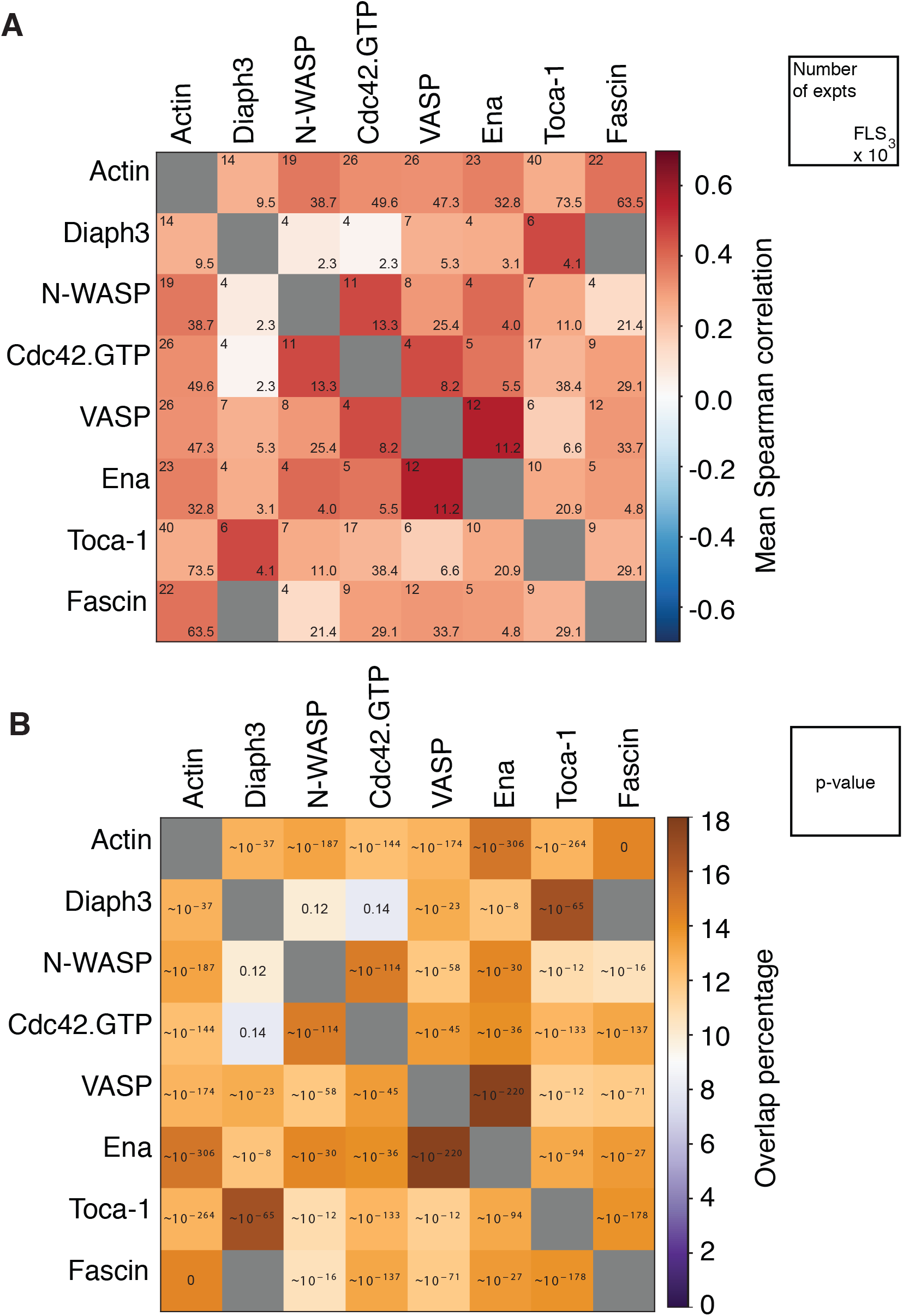
Heterogeneity and positive correlations between actin regulators at FLS tip complexes. (A) The correlation value matrix from Fig. 2B with numbers of experiments (upper left) and numbers of FLS (lower right, in thousands). (B) Percentage of FLS with two specific proteins enriched (above the 70th percentile in intensity), when both proteins are measured in the experiment. P-values (Cressie-Read power divergence test) for significant deviation from the 9% completely random overlap are given inside the boxes, zero indicates a level so small it is effectively zero.

**Fig. 2 - supplement 1.**
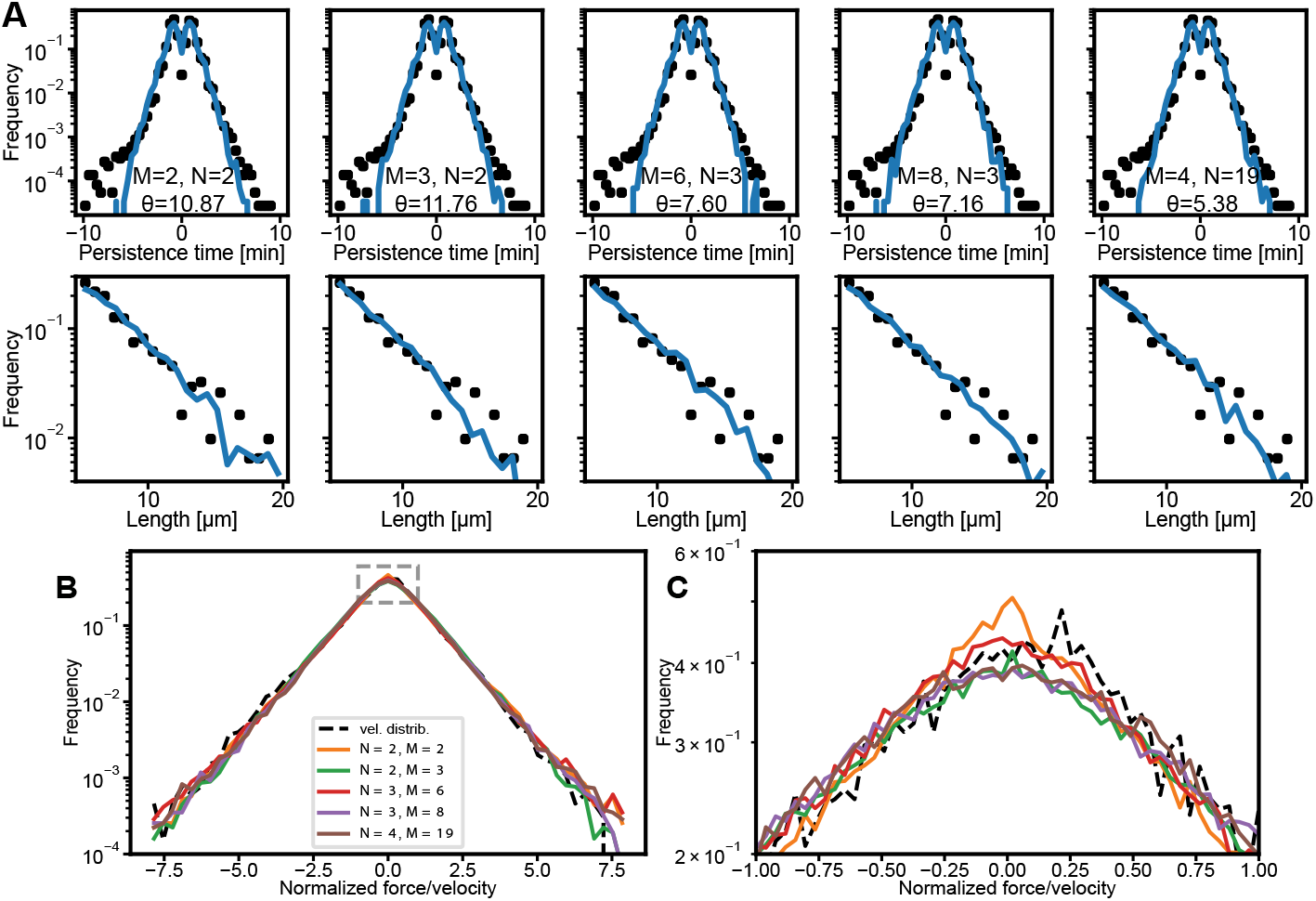
Theory fits to experimental data. (A) Predicted persistence time distribution (top row, blue solid lines) and length distributions (bottom row, blue solid lines) agree with the experimental distributions (black dots) for different values of M and N. Values for θ are given in min^−1^. (B) Force/velocity distributions for the five combinations highlighted by colored squares in Fig. 3E. The dashed line indicates the normalized experimental growth velocity. (C) An enlargement of the peak of the distributions indicated by the dashed rectangle in (B).

**Fig. 3 - supplement 1.**
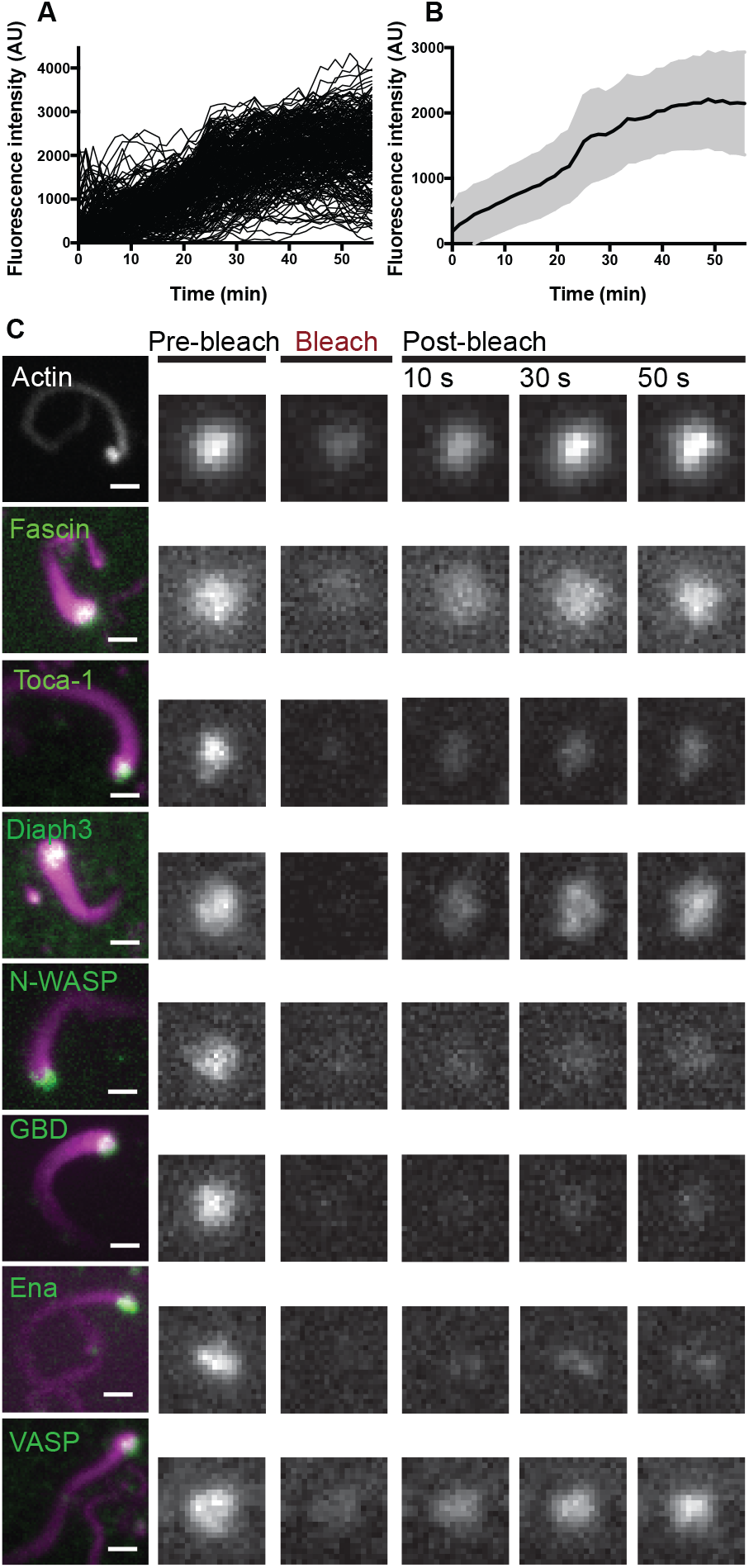
Actin intensity at FLS tips over time and examples of fluorescence recovery of actin regulatory proteins after photobleaching. (A) Individual timelapse traces of actin intensity in FLS showing stabilization at 30-40 mins (n=246) (B) Averaged traces of actin intensity, shaded area represents the standard deviation. (C) Example images actin regulatory protein localization to FLS, and fluorescence recovery after photobleaching at FLS tips (green: protein of interest, magenta AF647 actin). Scale bar = 2 *μ*m.

## Supplementary Information

### Mathematical Analysis

Like in the overdamped limit of Langevin equations (i.e. low Reynolds number), the velocity can simply be set equal to the force. We then assume that the force (and thus the growth velocity) acting on the growing FLS F is a function of the fluctuating concentrations of a set of proteins Ai with i = 1,…,K. We assume that the proteins obey independent Ornstein-Uhlenbeck relaxation processes

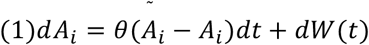

i.e. the protein concentrations fluctuate around a baseline concentration *Ã_i_*. The parameter *θ* > 0 determines the relaxation time scale of fluctuations. The Brownian process *W*(*t*) has zero mean and obeys the correlations 〈*W*(*t*)*W*(*s*)〉 = 2*θη*^2^*δ*(*t* − *s*) with the noise controlled by the parameter *η*. We take the difference between concentration and baseline as *X_i_* = *A_i_* − *Ã_i_*, hence the *X_i_* are normal distributed with zero mean in the long time limit.

One thus expects force to arise from a combination of protein concentration, as indicated by the fact that we see weak, but non-zero correlations between FLS lengths and many proteins. The most minimalistic and generic version of such a model is to assume that each type of protein is independent of each other (uncorrelated random variables), but that they act on force in either an additive or a multiplicative way. For instance, if two proteins *X*_1_ and *X*_2_ act on the force via a common given complex, the output would be expected to be multiplicative *F* = *X*_1_*X*_2_, while if they interact in an independent pathway, the output would be expected to be additive *F* = *X*_1_ + *X*_2_. In general, we can thus see the output force as a sum of *M* products of *N* fluctuating protein concentrations:

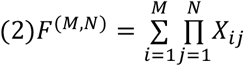

Strikingly an exact Laplacian force distribution can be achieved by the combination:

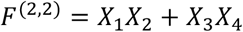

This can be shown analytically. Indeed the product of two standard normal distributions *Z* = *X*_1_*X*_2_ is distributed according to a modified Bessel function of the second kind:

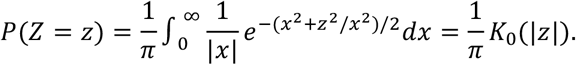

The sum of two products *W* = *Z*_1_ + *Z*_2_ is then distributed according to a Laplace distribution

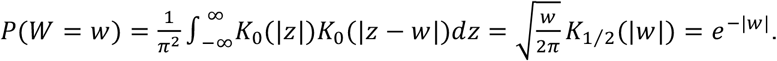

In order to simulate FLS growth, we generate *K* = *MN* artificial protein concentration traces by integrating (1) using the Euler-Maruyama method with a time step of one second and combine them according to equation (2) to generate the force. We then arrive at FLS length trajectories *L*(*t*) by temporal integration of the equation

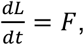

with the conditions *L*(*t*) ≥ 0 and *L*(*t* = 0) = 0.

### Biochemical connection

In order to make the connection between fluctuating protein concentrations and instantaneous force / growth velocity more tangible, we present a chemical reaction network consisting of 2M species Ai, which interact in pairs to form M complexes Bj. The complexes are subject to decay. The reaction network is thus:

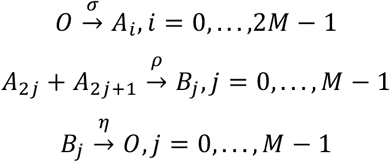

The mass action rate equations for the single species are then given by

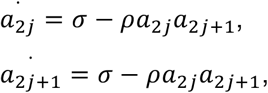

with the (degenerate) steady state solutions 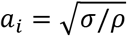. Therefore, we can write these concentrations in terms of the deviations (fluctuations) around the steady state:

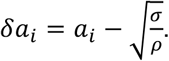

The rate equations for the complexes, expressed in terms of fluctuations are given by

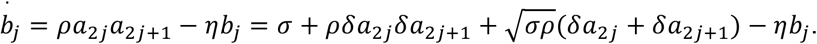

with the total velocity for an FLS reading:

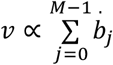

Thus, there are three contributions to the steady-state concentrations of the complex *b_j_* (and conversely to FLS speed). The first contribution arises from the steady rate of production *σ*, which dictates the average complex concentration. The product of such terms gives rise to a constant non-zero velocity of FLS growth. However, as steady state, FLS net average velocity is close to zero, because of depolymerization processes we have not modelled so far. Monomer availability for instance is a straightforward way to couple net polymerization and net depolymerization, ensuring that this first, non-zero average, contribution drops out. The second contribution *δa*_2*j*_*δa*_2*j*+1_ has already been discussed in the manuscript, and involves the product of Gaussian fluctuations, yielding generically exponential tails for *v* (and in the limit of two sums, an exact Laplace distribution). The third contribution 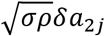 is a cross-term that involves the average concentration of one of the species and the fluctuation of the others, and is thus Gaussian distributed. However, our data is consistent with species *a_i_* with large variance compared to mean. In this limit, the third, Gaussian contribution becomes negligible and the global distribution for velocities approaches a Laplace distribution rapidly, with effectively the distribution controlled by 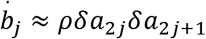. One should note that in this limit of large variance, the Ornstein-Uhlenbeck equations for the concentrations *a_i_* must be amended with reflective boundary conditions at zero to avoid negative concentrations (or alternatively, a steep potential around 0). Importantly, we tested that this still generically yielded Laplace-like distributions, for instance when combining regulators of growth vs. regulators of shrinkage, effectively generalizing the expression above to:

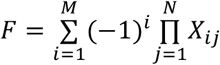

with each species *X_ij_* strictly positive and obeying the modified stochastic dynamics mentioned above (as mentioned above, we assume that there are feedback mechanisms to ensure that the sum of all averages is zero, yielding steady-state conditions for the force/velocity).

Finally, under the assumption that force / growth velocity generation is directly dependent on the concentrations of each of the regulatory complexes, we arrive at the sum of products rule mentioned in the main text:

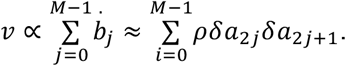

## Video Legends

**Video 1.**

Timelapse z-stack spinning disk confocal video of actin visualized with GFP-Utrophin CH domain probe at 20-30 min.

**Video 2.**

Maximum intensity projection of laser scanning confocal timelapse z-stacks of *enaGFP, en-Gal4; UAS-cd8mCherry* embryos, late stage 14. Representative video of 12 from 7 embryos.

**Video 3.**

Maximum intensity projection of laser scanning confocal timelapse z-stacks of *Scar/WAVENeonGreen en-Gal4; UAS-cd8mCherry* embryos, late stage 14. Representative video of 13 from 8 embryos.

**Video 4.**

Maximum intensity projection of laser scanning confocal timelapse z-stacks of *enaGFP, btl-Gal4; UAS-cd8mCherry* embryos, late stage 14. Representative video of 24 from 20 embryos.

**Video 5.**

Maximum intensity projection of laser scanning confocal timelapse z-stacks of *Scar/WAVENeonGreen; btl-Gal4 UAS-CAAXCherry* embryos, stage 15. Representative video of 9 from 6 embryos.

**Videos 6 and 7**

Maximum intensity projections of laser scanning confocal timelapse z-stacks of *GFP-fascin, en-Gal4; UAS-cd8mCherry* embryos, stage 15. Representative videos of 17 from 14 embryos.

**Data File – Figure 1 Panel B**

JavaScript Object Notation (JSON) file containing a list of segmented FLS time series data. Used to generate the average protein intensity time courses shown in Figure 1B and the temporal information in Figure 1 - supplement 3.

**Data File – Figure 1 Panel D, E and F**

Comma-separated table containing the data underlying the intensity and morphology correlation in Figure 1D, the relative FLS length data in Figure 1E and the FLS length distribution data in Figure 1F.

**Data File – Figure 1 supplement 4 Panel B**

Comma-separated table containing the number of local protein maxima detected with and without Latrunculin B, displayed in Figure 1S4 Panel B.

**Data File – Figure 2 Panels A and B**

Comma-separated table containing the time course information for the FLS segmentation of the high-timeresolution Utrophin-GFP timelapse movies. Used to generate Figure 2A and Figure 2B.

**Data File – Figure 3 Panel A**

Comma-separated table containing the FRAP time curve data used to generate Figure 3A.

**Data File – Figure 3 Panel B and C**

Comma-separated table containing the FRAP half-time and recovery analysis results shown in Figure 3B and Figure 3C.

**Data File – Figure 4 Panel A**

Comma-separated table containing the data used to generate the velocity-intensity correlation graphs shown in Figure 4A.

**Data File – Figure 4 Panels C, D and E**

Comma-separated table containing the Ena/VASP depletion FLS quantification data used to generate Figure 4C, Figure 4D and Figure 4E.

**Data File – Figure 5 Panels E and F**

Comma-separated table containing the Ena and Scar filopodia tip intensity quantification data shown in Figure 5E and Figure 5F.

**Data File – Figure 5 Panel G**

Comma-separated table containing length data for FLS, filopodia in *Drosophila* embryo dorsal closure leading edge cells and filopodia in *Drosophila* embryo tracheal cells, shown in Figure 5G.

**Data File – Figure 6 Panels B and D**

Comma-separated table containing segmented *Drosophila* embryo dorsal closure leading edge cell filopodia shaft fascin intensities with associated background intensities and length data. The intensity data is shown in Figure 6B, and the intensity-length scatter plot is shown in figure 6D.

**Data File – Figure 6 Panel C**

Comma-separated table containing FLS shaft fascin intensity to background ratios shown in Figure 6C.

**Data File – Figure 6 Panels E and F**

Comma-separated table containing segmented *Drosophila* embryo dorsal closure leading edge cell filopodia lengths for wild-type control, GFP-fascin knock-in, mutant fascin and fascin overexpression conditions, shown in Figure 6E and Figure 6F.

